# Metabolic dysfunction induced by high-fat diet modulates hematopoietic stem and myeloid progenitor cells in brown adipose tissue of mice

**DOI:** 10.1101/2021.03.02.433510

**Authors:** Kyle T Mincham, Kunjal Panchal, Prue H Hart, Robyn M Lucas, Martin Feelisch, Richard B Weller, Vance B Matthews, Deborah H Strickland, Shelley Gorman

**Affiliations:** Telethon Kids Institute, University of Western Australia, Perth, Australia; National Centre for Epidemiology and Population Health, Research School of Population Health, Australian National University, Canberra, Australian Capital Territory; Centre for Ophthalmology and Visual Science, University of Western Australia, Perth, Australia; Clinical and Experimental Sciences, Faculty of Medicine, University of Southampton, Southampton General Hospital, Southampton, United Kingdom; University of Edinburgh, MRC Centre for Inflammation Research, Edinburgh, Scotland; School of Biomedical Science - Royal Perth Hospital Unit, The University of Western Australia, Perth, Australia

**Keywords:** adiposity, bone marrow, brown adipose tissue, dendritic cells, high-fat diet, mice, metabolic dysfunction, myeloid cells, nitric oxide, stem cell, ultraviolet radiation

## Abstract

Brown adipose tissue (BAT) may be an important metabolic regulator of whole-body glucose. While important roles have been ascribed to macrophages in regulating metabolic functions in BAT, little known is known of the roles of other immune cells subsets, particularly dendritic cells (DCs). Eating a high fat diet may compromise the development of hematopoietic stem and progenitor cells (HSPC) – which give rise to DCs – in bone marrow, with less known of its effects in BAT. We have previously demonstrated that ongoing exposure to low-dose ultraviolet radiation (UVR) significantly reduced the ‘whitening’ effect of eating a high-fat diet upon interscapular (i)BAT of mice. Here, we examined whether this observation may be linked to changes in the phenotype of HSPC and myeloid-derived immune cells in iBAT and bone marrow of mice using 12-colour flow cytometry. Many HSPC subsets declined in both iBAT and bone marrow with increasing metabolic dysfunction. Conversely, with rising adiposity and metabolic dysfunction, conventional (c)DCs increased in both of these tissues. When compared to low-fat diet, consumption of high-fat diet significantly reduced proportions of myeloid, common myeloid and megakaryocyte-erythrocyte progenitors in iBAT, and short-term hematopoietic stem cells in bone marrow. In mice fed a high-fat diet, exposure to low-dose UVR significantly reduced proportions of cDCs in iBAT, independently of nitric oxide release from irradiated skin (blocked using the scavenger, cPTIO), but did not significantly modify HSPC subsets in either tissue. Further studies are needed to determine whether changes in these cell populations contribute towards metabolic dysfunction.

## Introduction

Excessive caloric intake contributes to the development of chronic diseases of metabolic dysfunction such as obesity and type-2 diabetes. These diseases incur large health and economic burdens in many regions of the world. Excessive caloric intake has systemic effects on multiple tissues including metabolically active sites, such as brown adipose tissue (BAT), an important regulator of whole-body glucose and temperature homeostasis. BAT has been detected in ∼7% of human adults^1^, with deposits occurring in the cervical, supraclavicular, axillary, and paravertebral regions^2^. These can vary significantly, ranging in mass from <1 to >150 g^2^. Factors further influencing BAT detection, activity and mass include: body mass index, gender, age and outdoor temperature^1, 2^. In addition, increased caloric intake may activate thermogenesis (heat production) in BAT through uncoupling-1 protein to help control glucose homeostasis and insulin sensitivity^3^. This process may be controlled by activities of immune cells.

Interactions between type-2 innate cells (lacking B/T cell receptors, producing type-2 cytokines), eosinophils and alternatively-activated (M2-type) macrophages promote the activation of BAT and browning of white adipose tissue (WAT) through type-2 cytokines such as interleukin-4 and −13 (reviewed^4^). Conversely, pro-inflammatory cytokines, such as tumour necrosis factor and interleukin-1ß, are produced by pro-inflammatory (M1-type) macrophages and may reduce uncoupling-1 protein expression and limit thermogenesis in BAT^4^. Other potentially important immune cells identified in BAT include regulatory T cells^5^ and B cells^6^. In many tissues, particularly those with more direct interactions with the external environment (e.g. lungs, intestines, skin), myeloid-derived dendritic cells (DCs) direct immune responses through their antigen-presenting capacity. In WAT, DCs may help maintain tissue homeostasis, with possible functions including: antigen sampling, mediator production, T cell memory, and cross-talk with adipocytes^7^. However, little is known about the phenotype and function of myeloid cells and DC in BAT.

Bone marrow is a rich site of hematopoietic stem and progenitor cells (HSPC) which give rise to immune cells of myeloid and lymphoid lineages. The consumption of a high-fat diet substantially modifies the architecture of bone marrow, increasing adiposity and restructuring bone^8, 9^, while also changing the composition and self-renewing capacity of progenitor cells^9-11^. In preclinical animal models, high-fat diet may decrease the proliferative capacity of HSPC in bone marrow^10, 11^, potentially impacting their ability to repopulate distant sites, including adipose tissues, and their capacity to modulate metabolic and immune responses. Feeding mice a high-fat diet for a ‘long-term’ (i.e. ≥8 weeks) may reduce stem cell proportions (Sca-1^+^CD45^-^) in “vascular stromal cells” (non-adipose cells) isolated from epididymal (visceral) WAT^12, 13^. In contrast, increased proliferative capacity of adipose stem and progenitor cells was observed at the same site after ‘short-term’ feeding (for 7 days) mice a high-fat diet^14^. These findings point towards time-dependent (i.e. length of exposure) effects of high-fat diet, and potential controversies in this research field. Furthermore, there may be important links between adipose tissue-resident stem and progenitor cell function and immunity, with IL-33 produced by progenitor cells located in epididymal and mesenchymal WAT depots required to control the activity of type-2 innate cells as well as adipose tissue expansion, and immune homeostasis^14^. It may be that similar pathways of control occur in other adipose tissue types, such as BAT. The specific effects of high-fat diet on HSPC in BAT are important to consider as these may differ between WAT and BAT, with HSPC subsets in BAT not well-characterised. Furthermore, there is variability in the responsiveness of HSPC from anatomically diverse depots of WAT^14^.

Here, we explored the associations between increasing adiposity and metabolic dysfunction induced by ‘long-term’ feeding mice a high-fat diet, and HSPC and myeloid cell proportions and numbers in interscapular (i)BAT and bone marrow. We used our recently established flow cytometry methods from similar studies done in lung and bone marrow^15^ to phenotype a hierarchy of HSPC and myeloid-derived cells, with a focus on DCs. This was done as a sub-study of now published experiments^16^, in which we investigated the effects of low dose ultraviolet radiation (UVR) on metabolic dysfunction in mice fed a high-fat diet, with a focus on iBAT^16^. In that study, we identified important metabolic benefits in iBAT, dependent on skin release of nitric oxide following exposure to UVR, including a reduced ‘whitening’ (i.e. WAT phenotype) effect of high-fat diet^16^. As (white) adipose tissue resident progenitor cells may have similar beneficial effects (to UVR) in preventing the expansion of adipose tissue^14^, we also report here effects of UVR on HSPC and myeloid cells in mice exposed to low dose UVR (and fed a high-fat diet), with and without further topical treatment with a nitric oxide scavenger (cPTIO). We first described the nature of observed associations between various measures of adiposity and metabolic dysfunction, and HSPC and myeloid cells in iBAT and bone marrow in the combined dataset. We then compared data between treatments, examining the direct effects of high-fat diet and ongoing exposure to low dose UVR, with or without topical treatment with cPTIO.

## Results

### Inverse linear correlation between iBAT mass and insulin sensitivity

We first considered the relationships among metabolic and adiposity outcomes within the total (combined) dataset, with data pooled together across all four treatments and each mouse considered as an individual datapoint (*n*=24). As expected, there were significant linear correlations between weight gain (throughout the 12-week treatment period), and final body weight (Figure 1a, r = 0.68, *P* = 0.0003) or gonadal (g)WAT mass (Figure 1b, r = 0.74, *P* < 0.0001). A positive correlation was observed between body weight and gWAT mass (Figure 1c, r = 0.49, *P* = 0.01) and insulin tolerance test area under the curve (Figure 1d, r = 0.50, *P* = 0.01). Similarly, there were significant positive linear correlations between iBAT mass and body weight (Figure 1e, r = 0.45, *P* = 0.03) or insulin tolerance test area under the curve (Figure 1f, r = 0.52, *P* = 0.009). No other significant correlations were observed amongst metabolic and adiposity outcomes in this dataset, in which fasting insulin, fasting glucose, and glucose tolerance test area under the curve were also considered, although non-significant correlations between fasting insulin and fasting glucose (r = 0.39, *P* = 0.06) or liver mass (r = 0.36, *P* = 0.09) were observed (data not shown).

**Figure 1.**
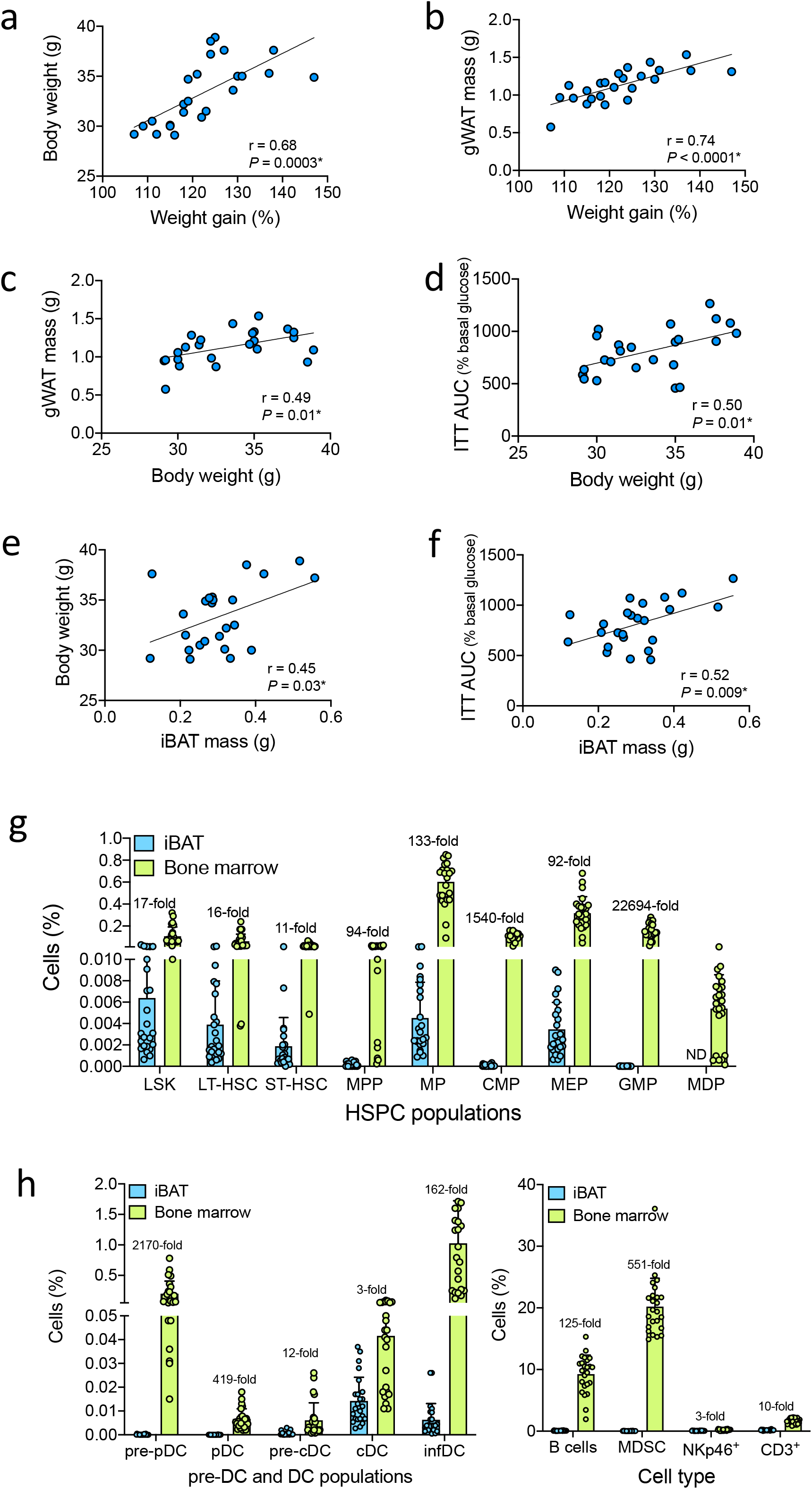
iBAT mass positively correlates with body weight and insulin tolerance test AUC with some HSPC and committeed myeloid cells were very infrequently or not detected in iBAT. Positive correlations between weight gain and (a) body weight or (b) gWAT mass, or, body weight and (c) gWAT mass or (d) ITT AUC (insulin tolerance test area under the curve), or, iBAT mass and (e) body weight or (f) ITT AUC were observed for data combined from 4 treatment groups for the experiment described in Figure 8. Two technical repeats of the same experiment were done with results combined, with three mice from each treatment per technical repeat combined for 6 mice per treatment. Data are shown for *n*=24 mice compared using Pearson correlation (r) test (\**P* < 0.05). In (g) and (h), the percentage of different HSPC, committed myeloid and other cells in iBAT (vascular stromal) cells and bone marrow cells collected at the end of the experiment are shown (ND=not detected). Fold differences in the percentages of each type between iBAT and bone marrow are shown, with data depicted as mean + SD. LSK=lineage^-^Sca-1^+^cKit^+^ cells; LT-HSC=long-term hematopoietic stem cell; ST-HSC=short-term hematopoietic stem cell; MPP=multipotent progenitor; MP=myeloid progenitor; CMP=common myeloid progenitor; MEP=megakaryocyte-erythrocyte progenitor; GMP=granulocyte-macrophage progenitor; MDP=macrophage-dendritic cell progenitor; pre-pDC=pre-plasmacytoid dendritic cell; pDC=plasmacytoid dendritic cell; pre-cDC=pre-conventional dendritic cell; cDC=conventional dendritic cell; infDC=inflammatory dendritic cell; MDSC=myeloid-derived suppressor cell.

### Some HSPC and myeloid cell types were infrequently or not detected in iBAT

We next compared the proportions of HSPC and committed myeloid cells between iBAT (total vascular stromal cells) and bone marrow (long bone cells) using our established flow cytometry methods^15^. Data were again pooled across all four treatments with each mouse considered separately (*n*=24). A limitation of our approach was that red blood cells were not lysed from the vascular stromal cell fraction isolated from iBAT during the cell isolation procedure, with initial optimisation trials suggesting that cell viability would be compromised when combined with the tissue digestion methodology. By contrast, red blood cells were lysed from cells isolated from the bone marrow as per our previously optimised methods^15^. There were significantly lower proportions of all HSPC in iBAT compared to bone marrow (Figure 1g, p≤0.0007, using multiple student’s *t-*tests (paired) with Bonferroni-Dunn correction for multiple tests). Indeed, some cell subsets were rarely or not detected in iBAT, including common myeloid progenitors (CMP), granulocyte-macrophage progenitors (GMP), macrophage-dendritic cell progenitors (MDP), pre-plasmacytoid DC or pre-conventional DC (Figure 1g, h; see also Supplementary table 1). Indeed, compared to bone marrow, there were far fewer myeloid progenitors (MP) that expressed CD34 and/or CD16/32 (Supplementary figure 1). The gating for CMP and GMP in iBAT was determined by our previous experience with tissue-specific profiling of these populations^15^, and that of seminal studies done in this field^17^. Increased proportions of all committed immune cell subsets were detected in bone marrow compared to iBAT; including plasmacytoid DCs, conventional DCs (cDC), inflammatory DCs, B cells and myeloid-derived suppressor cells (Figure 1h, p≤0.0005).

### Proportions of conventional DCs increased in iBAT with increasing adiposity

We then considered the relationships between proportions of HSPC and committed myeloid cells in iBAT, and metabolic outcomes. No significant relationships were observed between proportions of any HSPC subtype in iBAT and any metabolic outcome (Supplementary table 1). Positive correlations were observed for the proportions of cDC in iBAT and two markers of adiposity; body weight (Figure 2a, r = 0.41, *P* < 0.05), and iBAT mass (Figure 2b, r =0.41, *P* < 0.05; Supplementary table 1).

**Figure 2.**
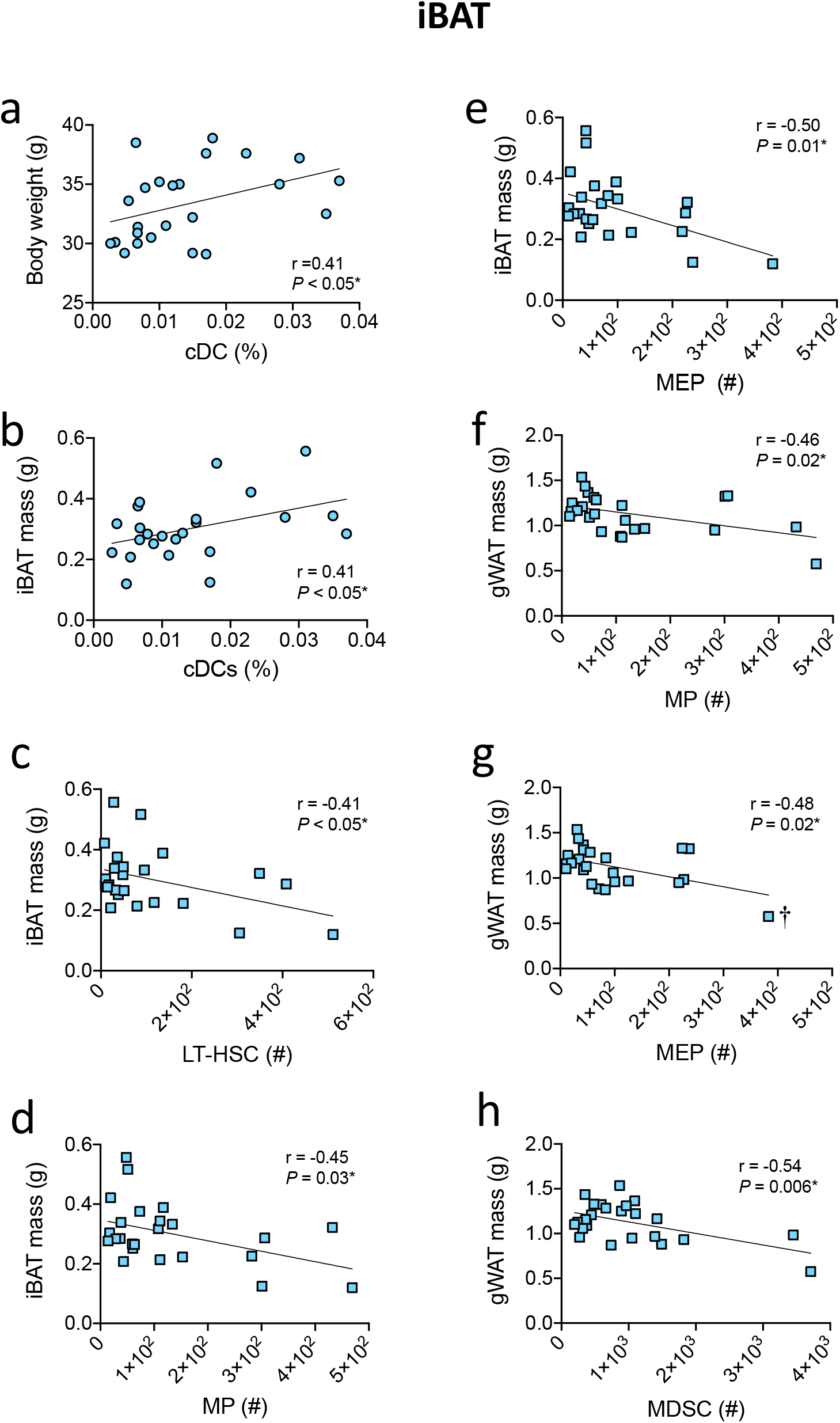
cDC increase in prevalence in iBAT with increasing adiposity, while some HSPC subsets decline in number as adipose tissue mass increases. Positive linear relationships were observed between proportions of conventional dendritic cells (cDC) and (a) body weight or (b) iBAT mass. Negative linear relationships were observed between numbers (per g of iBAT) of: (c) long-term hematopoietic stem cells (LT-HSC), (d) myeloid progenitors (MP), (e) megakaryocyte-erythrocyte progenitors (MEP) and iBAT mass; and, (f) MP, (g) MEP or (h) myeloid-derived suppressor cells (MDSC) and gBAT mass. Data were combined from 4 treatment groups of the experiment described in Figure 8 for data obtained from *n*=24 individual mice. Two technical repeats of the same experiment were done with results combined, with three mice from each treatment per technical repeat combined for 6 mice per treatment. Data were compared using Pearson correlation (r) tests (\**P* < 0.05), with no outliers identified (via ROUT test), although following the exclusion of a more extreme datapoint (†) in (g) changed the outcome of the correlation to non-significant (r = −0.20, *P* = 0.36).

### MP and MEP decreased in number in iBAT as iBAT or gWAT mass increased

We next considered the relationships between numbers (per g of tissue) of HSPC and committed myeloid cells in iBAT, and, metabolic outcomes (Supplementary table 2). Significant positive linear relationships between numbers of long-term hematopoetic stem cells (LT-HSC; Figure 2c, r = −0.41, *P* < 0.05), MP (Figure 2d, r = −0.45, *P* = 0.03), and megakaryocyte-erythrocyte progenitor (MEP; Figure 2e, r = −0.50, *P* = 0.01) cells and iBAT mass were observed. Similar relationships were observed between these HSPC subsets and gWAT mass (LT-HSC; data not shown, r = −0.38, *P* = 0.07; MP; Figure 2f, r = −0.46, *P* = 0.02; MEP; Figure 2g, r = −0.48, *P* = 0.02). There was also a significant inverse relationship between multipotent progenitor (MPP) numbers in iBAT and fasting insulin (Supplementary table 2; r = −0.41, *P* = 0.047). These data suggest that with increasing iBAT or gWAT mass, MP and MEP HSPC populations decline in number in iBAT. A negative relationship was also observed between myeloid-derived suppressor cell numbers in iBAT and gWAT mass (Figure 2h, r = −0.54, *P* = 0.006). Significant negative correlations were similarly observed between both percentage (r = −0.42, *P* = 0.041) or numbers (r = −0.59, *P* = 0.002) of neutrophils in iBAT and gWAT mass (see Supplementary tables 1, 2). However, significant variation was observed, with outlier datapoints identified (via ROUT test) within the neutrophil percentage (*n* = 1) and number (*n* = 2) datasets. When these outliers were excluded, the correlations of neutrophil percentages (r = −0.14, *P* = 0.53) or numbers (r = −0.36, *P* = 0.11) with gWAT mass were no longer significant.

### HSPC declined in bone marrow with increasing metabolic dysfunction

In bone marrow, significant inverse correlations were observed between the proportions of LT-HSC (Figure 3a, r = −0.44, *P* = 0.03), MPP (Figure 3b, r = −0.64, *P* = 0.0008), and MDP (Figure 3c, r = −0.66, *P* = 0.0005) and fasting insulin (Supplementary table 3). Similar significant inverse relationships were observed for numbers of MPP (Supplementary table 4, Supplementary figure 2a, r = −0.55, *P* = 0.005) or MDP (Supplementary figure 2b, r = −0.55, *P* = 0.005) in bone marrow and fasting insulin. A significant negative correlation was observed between proportions of short-term hematopoietic stem cells (ST-HSC) and iBAT mass (Figure 3d, r = −0.42, *P* = 0.04). When considering fasting glucose, inverse relationships between both proportions (Figure 3e, r = −0.52, *P* = 0.009) and numbers (Supplementary figure 2c; r = −0.72, *P* < 0.0001) of MEP were observed. Significant negative correlations were also observed for lineage^-^Sca-1^+^c-Kit^+^ cells, MP, and CMP numbers in bone marrow and fasting glucose (Supplementary figure 2d-f; −0.62 < r ≤ −0.43, *P* ≤ 0.03).

**Figure 3.**
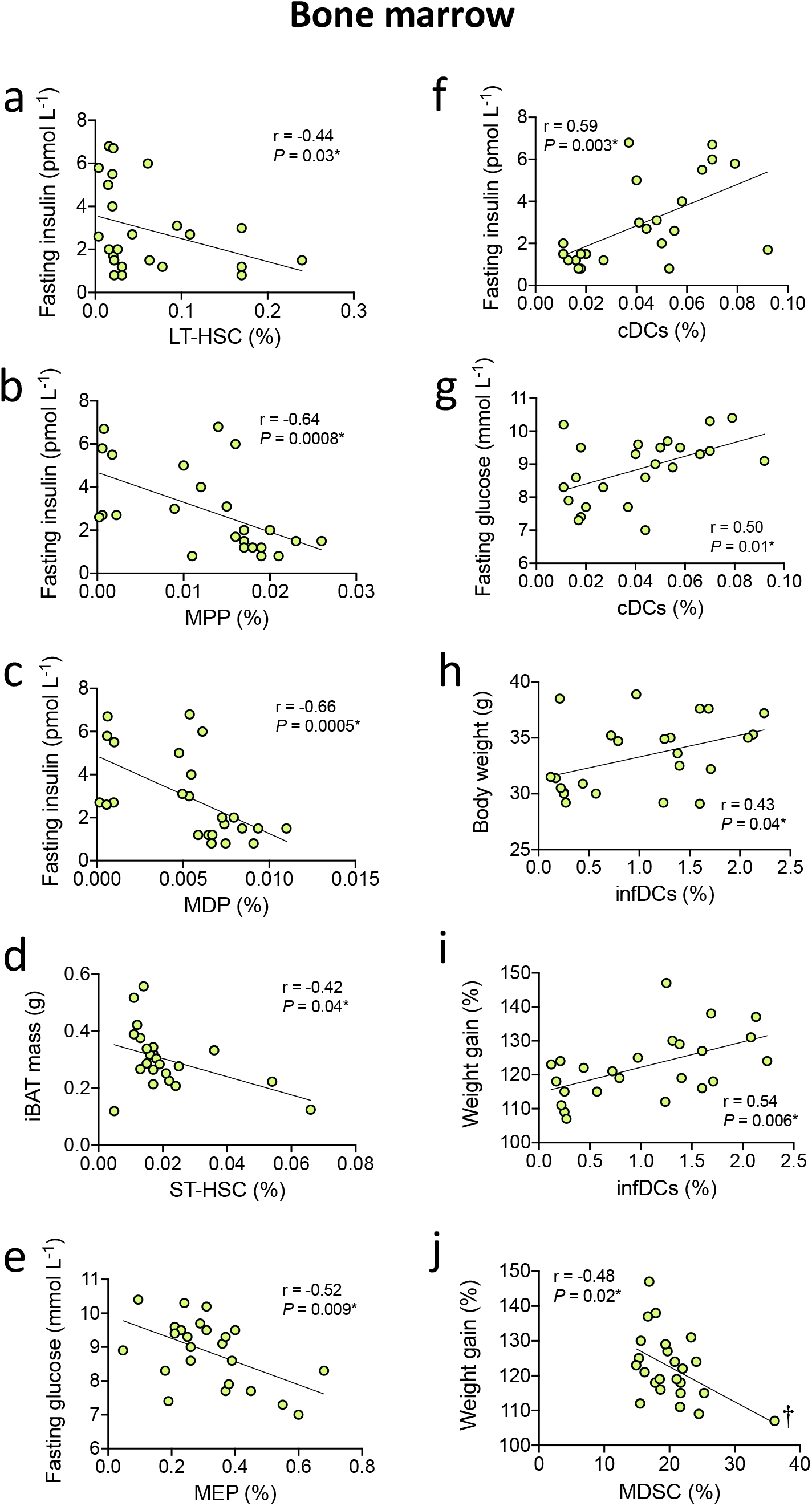
Increasing metabolic dysfunction and adiposity was associated with reductions in the proportions of some HSPC subsets, and conversely (mainly) increases in some DCs subtypes in bone marrow. Inverse linear relationships between percentages of: (a) long-term hematopoietic stem cells (LT-HSC), (b) multipotent progenitors (MPP), or (c) macrophage-dendritic cell progenitor (MDP), and fasting insulin were observed. (d) Inverse relationship between percentage of short-term hematopoietic stem cells (ST-HSC) and iBAT mass. (e) Inverse linear relationship between percentage of megakaryocyte-erythrocyte progenitors (MEP) and fasting insulin. Positive linear relationships observed between (f) fasting insulin or (g) glucose levels and conventional dendritic cells (cDC) proportions in bone marrow. (h) Positive linear relationships were also observed for inflammatory DCs (infDCs) proportions in bone marrow, and (h) body weight, or (i) weight gain. (j) An inverse linear relationship was observed between proportions of MDSC and weight gain. Data were combined from 4 treatment groups of the experiment described in Figure 8 for data obtained from *n*=24 individual mice. Two technical repeats of the same experiment were done with results combined, with three mice from each treatment per technical repeat combined for 6 mice per treatment. All data were compared using Pearson correlation (r) test (\**P* < 0.05), except for fasting insulin data, which were compared using a Spearman correlation test as these data were not normally distributed. No outliers were identified (via ROUT test), although following exclusion of one more extreme datapoint (†) in (j) changed the outcome of the correlation to non-significant (r = −0.36, *P* = 0.09).

### cDCs and inflammatory DCs increased in bone marrow with increasing metabolic dysfunction

Positive relationships between proportions (Figure 3f, r = 0.59, *P* = 0.003) or numbers (Supplementary figure 2g, r = 0.45, *P* = 0.03) of cDC and fasting insulin were observed in bone marrow. Similar positive relationships were observed between the proportions of: cDC and fasting glucose (Figure 3g, r = 0.50, *P* = 0.01); and, inflammatory DCs and body weight (Figure 3h, r = 0.43, *P* = 0.04), weight gain (Figure 3i, r = 0.54, *P* = 0.006) or gWAT mass (Supplementary figure 2h, r = 0.50, *P* = 0.01). Significant negative relationships were observed between proportions of myeloid-derived suppressor cell in bone marrow and weight gain (Figure 3j r = −0.48, *P* = 0.02), or gWAT mass (Supplementary table 3: r = −0.50, *P* = 0.01). The same negative relationships were also observed for numbers of: pre-conventional DC and weight gain (Supplementary figure 2i, r = −0.45, *P* = 0.03); plasmacytoid DCs and insulin tolerance test area under the curve (Supplementary figure 2j, r = −0.45, *P* = 0.03); and, NKp46^+^ cells and fasting insulin (Supplementary figure 2k, r = −0.44, *P* = 0.03).

### Some significant correlations may have been driven by a single datapoint

For some significant correlations above, while no significant outlier data-points were identified through ROUT analysis, it was possible that these relationships were driven by a ‘more extreme’ single datapoint. When excluded, the significant relationships initially observed in Figure 2g (numbers of MEP in iBAT and gWAT), Figure 3j (proportions of myeloid-derived suppressor cells in bone marrow and weight gain (or gWAT, Supplementary table 3), and Supplementary figure 2 (d, f, i, k; numbers in bone marrow of lineage^-^Sca-1^+^c-Kit^+^ cells, or CMP and fasting glucose, or pre-conventional DC and weight gain, or NKp46^+^ cells and fasting insulin, respectively) became non-significant. The ‘more extreme’ datapoints in Supplementary figure 2(d, f, i, k) were acquired from the same bone marrow specimen.

### Ongoing exposure to low dose UVR reduced iBAT mass in mice fed a high-fat diet in this sub-study

We then considered the effects of each treatment separately on adiposity and metabolic outcomes (Figure 4). High-fat diet (alone) significantly increased body weight (Figure 4a) and gWAT mass (Figure 4d) measured at the end of the experiment, while significantly less iBAT mass was observed in mice fed a high-fat diet with ongoing exposure to low dose UVR (Figure 4f). These findings are largely consistent with those previously reported^16^, and as a sub-study of the original, were likely not sufficiently powered to demonstrate the effects of high-fat diet and other treatments on some metabolic outcomes. Even so, these findings suggest that iBAT may be a tissue that is relatively sensitive to the effects of low dose UVR.

**Figure 4.**
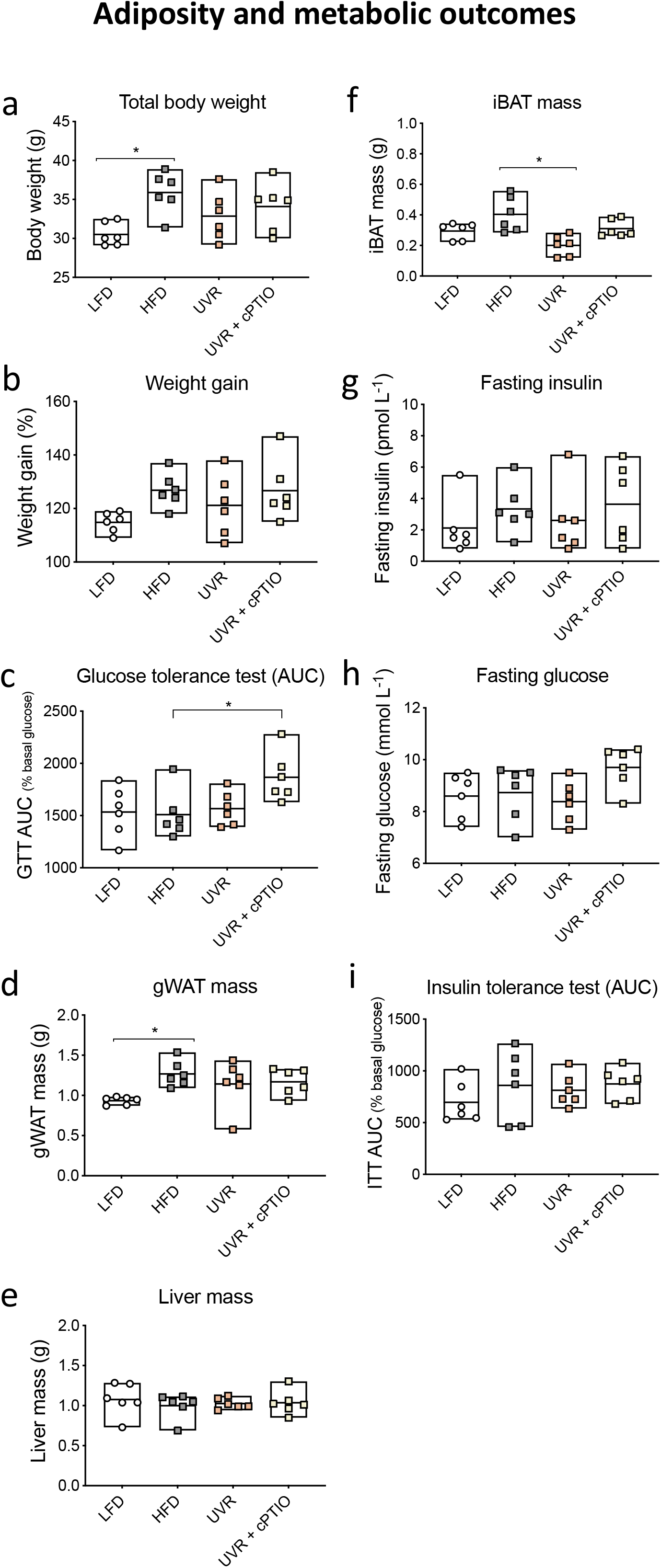
Ongoing exposure to low dose UVR reduced iBAT mass in mice fed a high-fat diet. Data are shown for mice for which HSPC and myeloid cells were characterised in iBAT and bone marrow, for 6 mice/treatment with the 4 treatment groups described in Figure 8. Two technical repeats of the same experiment were done with results combined, with three mice from each treatment per technical repeat combined for 6 mice per treatment. In (a) body weight, and (b) body weight gain are reported for data collected at the end of the experiment. In, (c) glucose tolerance test area under the curve (GTT AUC), performed at week 10. In (d-f), gonadal white adipose tissue (gWAT), liver and interscapular brown adipose tissue (iBAT) mass, collected from mice at the end of the experiment. In (g), fasting insulin (week 9), (h) fasting glucose (week 10) and (i) insulin tolerance test area under the curve (ITT AUC, week 11). Data were compared using ANOVA with Tukey’s post-hoc (if normally distributed) or Kruskal-Wallis test with Dunn’s post-hoc (if not normally distributed) (\**P* < 0.05). Data are shown for individual mice, with the mean (mid-line in box) and range (outer limits of box) depicted for each treatment.

### High-fat diet reduced some HSPC subsets in iBAT

Potential treatment-specific effects on HSPC and myeloid cells in iBAT and bone marrow were then examined. Proportions of MP, CMP and MEP in iBAT, and ST-HSC in bone marrow were significantly lower in mice fed a high-fat diet, compared to those fed the low-fat diet (Figure 5). When considering numbers of vascular stromal cells (non-adipose cells) in iBAT, high-fat diet significantly reduced the total number of these cells isolated per g of iBAT (Figure 6a), as well as numbers of lineage^-^Sca-1^+^c-Kit^+^ cells, LT-HSC, ST-HSC, MP, CMP and MEP (Figure 6b-g). There was no significant difference in the total cellularity (number of cells isolated) or numbers of specific HSPC subsets isolated from bone marrow between treatments (Figure 6h-n). Exposure to UVR (with or without cPTIO) did not consistently (or significantly) reverse the effects of high-fat diet on HSPC in iBAT or bone marrow, although for most populations there was no difference between mice fed low-fat diet and those exposed to UVR (Figure 5, 6). These findings suggest that high-fat diet significantly reduced HSPC populations, particularly in iBAT, and a lack of further effect of UVR was probably due to sample size limitations.

**Figure 5.**
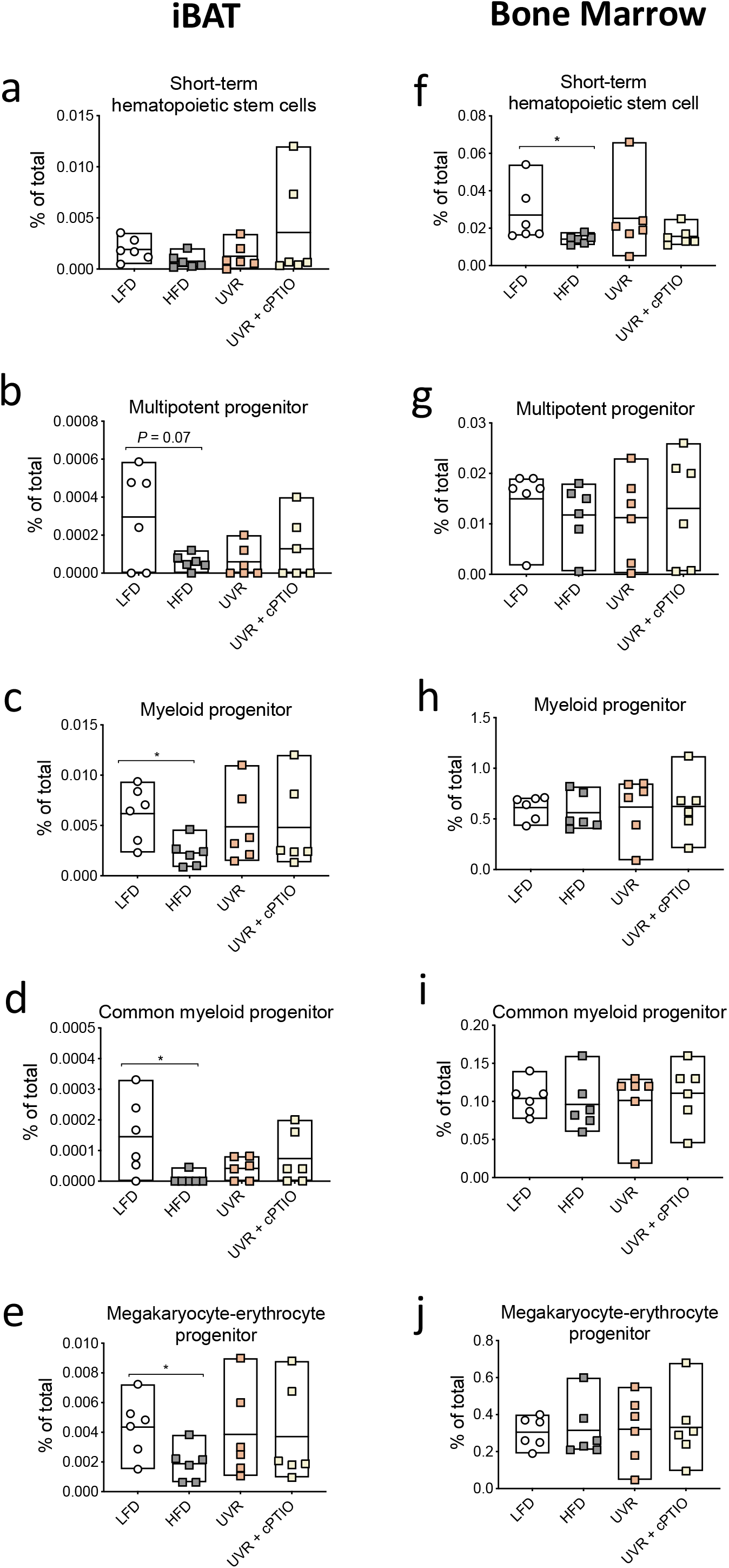
High-fat diet reduced proportions of someHSPC subsets in iBAT. Data are shown for iBAT (a-e) and bone marrow (f-j) collected from *n*=6 mice/treatment for the experiment described in Figure 8. Two technical repeats of the same experiment were done with results combined, with three mice from each treatment per technical repeat combined for 6 mice per treatment. The proportions of various HSPC in iBAT (% of total vascular stromal cells) and bone marrow are shown, including: (a, f) short-term hematopoietic stem cells (ST-HSC); (b, g) multipotent progenitor (MPP); (c, h) myeloid progenitor (MP); (c, i) common myeloid progenitor (CMP); and, (e, j) megakaryocyte-erythrocyte progenitor (MEP). Data were compared using ANOVA with Tukey’s post-hoc test (\**P* < 0.05). Data are shown for individual mice, with the mean (mid-line in box) and range (outer limits of box) depicted for each treatment.

**Figure 6.**
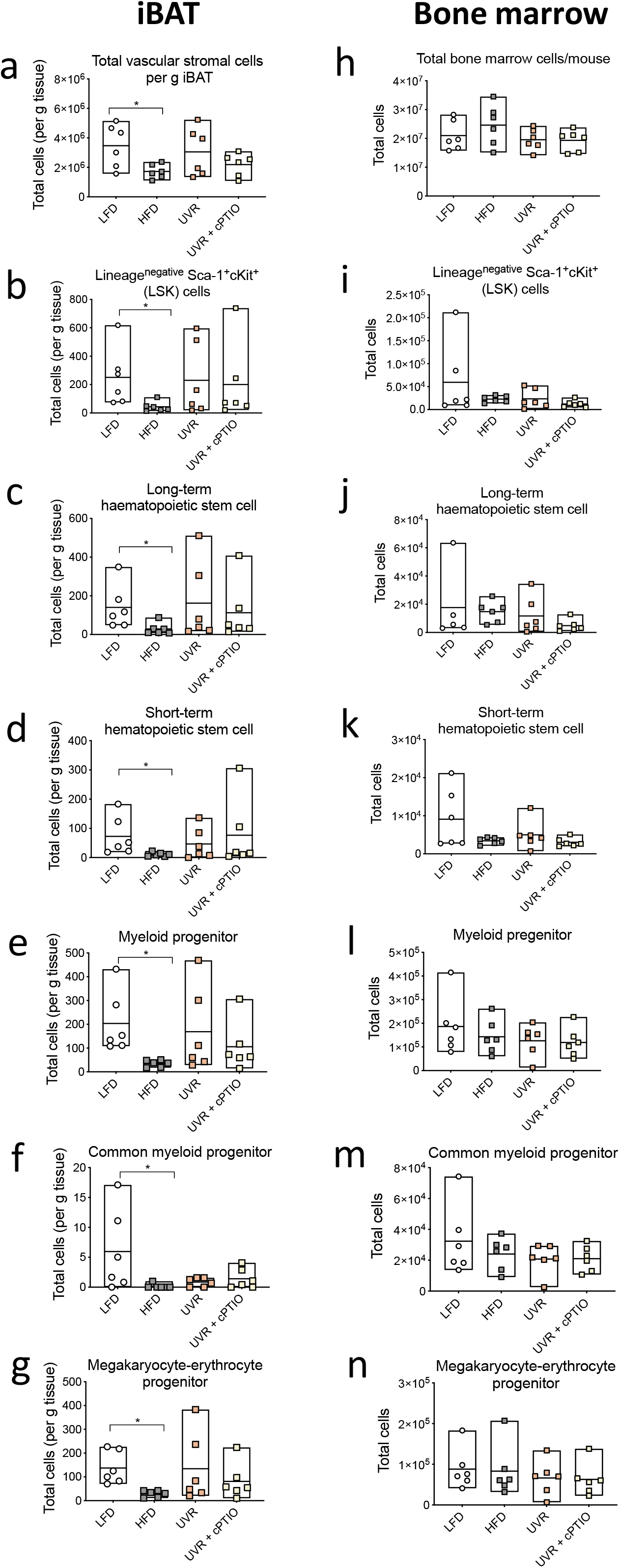
High-fat diet reduced HSPC numbers in iBAT but not bone marrow. Data are shown for iBAT collected from *n*=6 mice/treatment for the experiment described in Figure 8. Two technical repeats of the same experiment were done with results combined, with three mice from each treatment per technical repeat combined for 6 mice per treatment. The total number of vascular stromal (non-adipose) cells per g iBAT is shown in (a), and bone marrow cells in (h). Numbers of various HSPC (per g for iBAT) are shown, including; (b, i) lineage^-^ Sca-1^+^cKit^+^ cells (LSK), (c, j) long-term hematopoietic stem cells (LT-HSC), (d, k) short-term hematopoietic stem cells (ST-HSC), (e, l) myeloid progenitors (MP), (f, m) common myeloid progenitors (CMP) and (g, n) megakaryocyte-erythrocyte progenitors (MEP). Data were compared using ANOVA with Tukey’s post-hoc test (\**P* < 0.05). Data are shown for individual mice, with the mean (mid-line in box) and range (outer limits of box) depicted for each treatment.

### UVR reduced cDC in iBAT but not bone marrow

We then considered the effects of diet or exposure to UVR on committed myeloid populations in iBAT and bone marrow. Exposure to UVR significantly reduced proportions of cDC (but not numbers) in iBAT of mice fed the high-fat diet (Figure 7a, b). There was no significant difference in surface expression of MHC class II (IA/IE) on cDC isolated from iBAT of mice fed the high-fat diet compared to those fed the low-fat diet (*P* = 0.07, via unpaired student’s t-test, Figure 7c). The number of B cells in iBAT was lower in mice fed the high-fat diet compared to low-fat diet (Figure 7d). Finally, there was no effect of high-fat diet (with or without exposure to low dose UVR) on the proportions or numbers of any other committed cell subset identified in bone marrow (Figure 7g-l and data not shown). These data suggest that high-fat diet reduced B cell numbers, while exposure to UVR significantly diminished proportions of cDC in iBAT, but not bone marrow.

**Figure 7.**
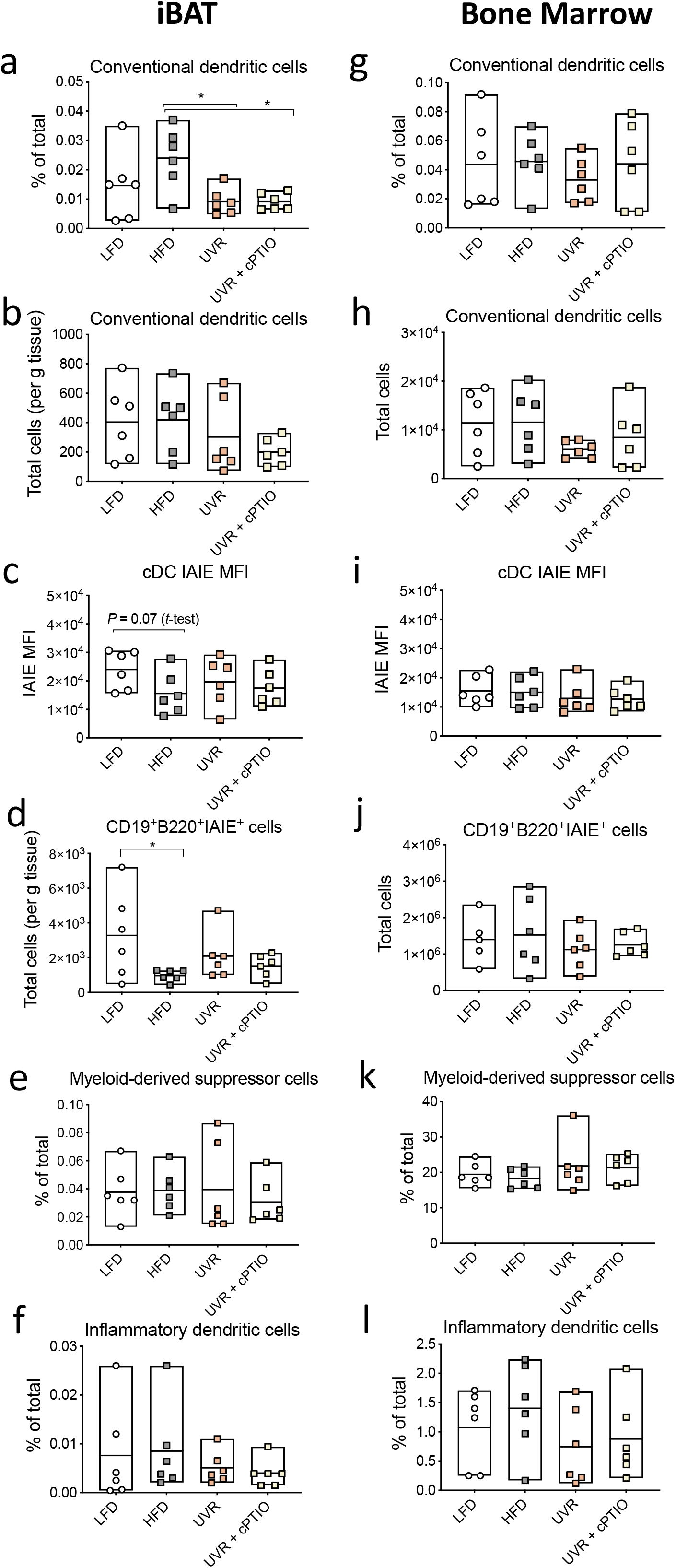
UVR reduced cDC proportions in iBAT. Data are shown for iBAT (a-f) and bone marrow (g-l) collected from *n*=6 mice/treatment for the experiment described in Figure 8. Two technical repeats of the same experiment were done with results combined, with three mice from each treatment per technical repeat combined for 6 mice per treatment. The proportions and numbers (respectively) of conventional DCs (cDC) in iBAT (a, b) and bone marrow (g, h) are shown. In (c) and (i), the mean fluorescent index (MFI) of MHC class II (IAIE) on cDC from iBAT and bone marrow (respectively). In (d) and (j), the numbers of CD19^+^B220^+^IAIE^+^ B cells iBAT and bone marrow (respectively). In (e) and (k), the proportions of myeloid-derived suppressor cells in iBAT and bone marrow (respectively). In (f) and (l), the proportions of inflammatory DCs in iBAT and bone marrow (respectively). Data were compared using ANOVA with Tukey’s post-hoc test (\**P* < 0.05). Data are shown for individual mice, with the mean (mid-line in box) and range (outer limits of box) depicted for each treatment.

## Discussion

Here, we phenotyped a hierarchy of HSPC and myeloid cells in iBAT and bone marrow, and analysed their associations with metabolic dysfunction, as well as the effects of high-fat diet and exposure to low dose UVR. Some cells were very infrequently or not detected in iBAT, including CMP, GMP, MDP, pre-plasmacytoid DC, and pre-conventional DC subsets. This may suggest that iBAT is not a likely niche for haematopoiesis. With increased metabolic dysfunction, HSPC significantly declined in both iBAT (LT-HSC, MPP, MP, MEP numbers) and bone marrow (LT-HSC, ST-HSC, MPP, MP, MDP, MEP numbers and/or proportions). Some of these effects could be explained by the effects of high-fat diet, with significantly lower LT-HSC, MP and MEP numbers observed in iBAT, and ST-HSC proportions in bone marrow (compared to mice fed a high-fat diet). Further studies are needed to determine whether eating a high-fat diet changes the way HSPC from bone marrow seed peripheral tissue (e.g. effects on differentiation, extravasation into circulation, apoptosis or other), and/or has direct effect on HSPC located in bone marrow, adipose tissues and other sites. High-fat diet also reduced B cell (CD19^+^B220^+^IA/IE^+^) numbers in iBAT compared to the low-fat diet. The impact of this finding is uncertain with other studies suggesting that B cells may participate in obesity-induced insulin resistance (reviewed in^18^). In iBAT, increased proportions of cDC were observed with increasing metabolic dysfunction, and in mice fed a high-fat diet these cells were significantly reduced with exposure to low dose UVR. As explored further below, this observation could be due to anti-inflammatory effects of UVR.

Other researchers have also observed significant reductions in ST-HSCs and suppression of progenitor populations in bone marrow with high-fat diet feeding^10, 11^. Similar observations have been made in WAT depots^12, 13^. However, an opposite effect has been observed in other studies where high-fat diet expanded myeloid progenitors (e.g. GMP^19, 20^) in bone marrow. These divergent observations could be explained by different markers and gating strategies used to define HSPC to those used here, with some previous studies^19, 20^ not using IL-7Rα^-^ and/or CD34 to define some important subsets (e.g. GMP)^17^. Hermetet *et al*. (2019) reported that high-fat diet disturbed the maintenance of HSPC in bone marrow, by impairing TGF-β-signalling in lipid rafts^11^. Our findings of fewer progenitors in bone marrow and iBAT in mice fed a high-fat diet suggest similar functional defects are induced by the accumulation of lipids in these tissues.

Through performing correlation analyses on the dataset as a whole, we also observed other potentially novel correlations between HSPC and myeloid subsets and metabolic dysfunction, particularly in the bone marrow for significant inverse relationships between LT-HSC, MPP, MP, MDP, MEP, and cDC subsets with fasting glucose and/or insulin. These findings point to potential links between glucose derangement and hormonal effects (e.g. via insulin), and haematopoiesis in bone marrow. However, while this approach allowed for consideration of variability in the measured metabolic outputs across all treatments, the causal direction of these significant relationships could not be determined. Even so, insulin signalling may control metabolic programming of stem cells in bone marrow^21^. We examined both cell proportions and cell numbers in these tissues for a more complete phenotyping of HSPC and myeloid cells. In iBAT, comparing the proportions of subsets within the total visceral stromal cell fraction was done to consider changes that were independent of size of iBAT, while changes in cell numbers per g tissue accounted for relative differences occurring due to expanding tissue mass (i.e. via deposition of fat). For some subsets (i.e. MPP, MDP) in bone marrow, significant correlations (r ≤ −0.55, *P* ≤ 0.005) were observed for both proportions and numbers of cells with fasting insulin that may be particularly worthy of future investigation.

We observed that ongoing exposure of mice twice-a-week to low dose UVR significantly reduced cDC in iBAT, independent of nitric oxide release from skin. We have previously seen both nitric oxide-dependent and independent benefits of UVR in preventing metabolic dysfunction in mice fed a high-fat diet^16^. UVR-induced nitric oxide limited the ‘whitening’ of iBAT (emergence of a WAT phenotype) and hepatic steatosis in mice fed a high-fat diet^16, 22, 23^. Nitric oxide released from skin did not mediate other changes induced by UVR in iBAT, although UVR exposure modulated the expression of mRNAs central to BAT function: increasing *Dio2, Glut4* and *Fatp2* mRNAs; and, reducing *Tnf, Atgl*, and *FasN* mRNAs^16^. Together, the reductions of *Tnf* mRNA levels and cDC proportions suggest that UVR has anti-inflammatory effects in iBAT that are independent of UVR-induced nitric oxide. TNF activates cDCs, promoting their migration into inflamed tissue and draining lymph nodes, and subsequent capacity to secrete pro-inflammatory cytokines^24-26^. Importantly, DCs in adipose tissue may contribute towards obesity-induced insulin resistance and inflammation^18^. Further experiments are needed to demonstrate if there are functional links between reductions in TNF levels and cDC in iBAT of mice exposed to UVR. Other studies indicate that UVR exposure can mediate functional changes in DCs derived from myeloid progenitors in bone marrow, including enhanced glycolytic flux, reduced migration and impaired capacity to prime immune responses^27-30^.

The ‘whitening’ of BAT which occurs in mice fed a high-fat diet is functionally linked to reduced vascularisation^31^. High-fat diet also increases iBAT mass^16^, which we show here to be positively correlated with markers of adiposity (body weight; r = 0.45, *P* = 0.03) and insulin insensitivity (i.e. insulin tolerance test area under the curve; r = 0.52, *P* = 0.009). We also observed reductions in MEP numbers in iBAT with increasing iBAT mass (r = −0.50, *P* = 0.01), and similar reductions in MEP proportions/numbers in bone marrow with increasing fasting glucose (r ≤-0.52, *P* ≤ 0.009). This finding was likely induced by mice eating a high-fat diet, with proportions and numbers of MEP also significantly reduced in iBAT of mice fed a high-fat diet (compared to low-fat diet). We hypothesise that these findings may be linked to the whitening effects of high-fat diet. Indeed, one cell type that differentiates from MEP is the erythrocyte and reductions in this cell type could contribute towards the ‘vascular rarefaction’ (reduced density of the microvascular network) that occurs during whitening of iBAT.

One reason why iBAT mass was reduced in mice exposed to UVR was due to its capacity to limit the whitening effect of eating a high-fat diet^16^. It is uncertain whether this observation was due to a change in the number of iBAT cells and/or their size, although whitening is likely linked to increased free fatty acid uptake and expansion of brown adipocytes. When considering total vascular stromal cells, these were reduced per g of iBAT in mice fed a high-fat diet compared to low-fat diet, which was not observed in mice exposed to UVR. These changes appeared to be independent of both numbers (per g tissue) or percentages of cDCs observed in iBAT. Further studies are needed to identify which cell types were increasing when cDCs were decreasing (in percentage); however, this could be difficult as cDCs were quite low in both number (<1000/g) and percentage (<0.1% of total vascular stromal cells) in iBAT.

In conclusion, we report here some evidence for inhibitory effects of consuming a high-fat diet on HSPC phenotypes in iBAT and bone marrow of experimental mice. Exposure to UVR significantly reduced cDCs in iBAT, independent of skin release of nitric oxide, which may be due to anti-inflammatory effects of UVR. A strength of this study was the design of these analyses as a sub-study of a larger experiment, using tissue from mice with metabolic outcomes already measured, and reducing the need for additional experimental animals. Important limitations include the relatively small sample size within each treatment group, and the use of an ‘anxious’ mouse strain (FVB/NJ) that may have different susceptibility to the effects of high-fat diet than other ‘more susceptible’ strains (e.g. C57Bl/6J)^32^. In addition, we did not correct for the multiple correlation tests performed, to avoid excluding possible false negatives that could be worthy of investigation in future studies. Further work is needed to reproduce these findings, considering important factors such as age and gender, and better define whether the metabolic benefits of UVR^16^ are related to changes in cDC proportions in iBAT, if functional links between MEP, vascularisation and whitening of BAT exist, and how findings here in mice are relevant to human biology. While obesity is associated with changes in the cell composition of bone marrow, including HSPC^33^, little is known of its effects on stem cells in human BAT.

## Methods

### Mice

As previously described^16^, this experiment was performed according to ethical guidelines of National Health & Medical and Research Council (Australia) with approval from the Telethon Kids Institute Animal Ethics Committee (AEC#315). Findings are reported according to ARRIVE (Animal Research: Reporting *In Vivo* Experiments) guidelines. Tg(*Ucp1*-luc2,-tdTomato)1Kajim/J transgenic mice (FVB/NJ background strain), also known as Thermomouse^34^, were obtained from the Jackson Laboratory (stock no. 026690; Bar Harbor, ME, USA) and housed and bred in the Bioresources Centre of the Telethon Kids Institute (Subiaco, Western Australia). Specific-pathogen-free experimental male mice were housed individually, which was an ethical requirement stipulated by the ethics committee due to excessive fighting exhibited by this mouse strain and sex. Mice exhibiting excessive anxiety (‘circling’ behaviour prior to 8 weeks of age) were excluded. From 4 weeks of age, mice were housed in temperature-controlled rooms (23.0 ± 0.9 °C, mean ± SD) under Perspex-filtered fluorescent lighting, with a normal 12 h light/dark cycle (lights on at 06:00 AM), with chipped Aspen bedding, tissues, crinkled paper and PVC piping for nesting in filter-topped cages. The fluorescent lighting does not emit any detectable UVR, as measured using a UV radiometer (UVX Digital Radiometer, Ultraviolet Products Inc., Upland, USA). Mice had access to food and water *ad libitum*.

### Experiment design

The findings reported here form a sub-study performed within a published experiment^16^, in which there were initially 20 mice per treatment. The choice for performing this sub-study was because of the opportunity and convenience of sampling, to enable tissue sharing to ‘reduce’ animal use, and to allow for comparison with our published data from the same experiments^16^. Mice were initially randomly allocated into treatments, with body weights compared (via one-way ANOVA) to confirm no baseline differences in body weight. In this sub-study, analyses of bone marrow and iBAT vascular stromal (non-adipose) cell populations and corresponding metabolic measures for individual mice were performed on a random selection of 6 mice per treatment (for all outcomes). This number of mice per group was selected based on prior experience of the number of samples that could be feasibly processed by researchers involved in data collection, considering also that cells from iBAT and bone marrow were to be individually phenotyped on a per mouse basis. Findings reported here were from two technical repeats of the same experiment (done in two ‘batches’) with results combined, as previously described^16^. Each technical repeat was initiated a week apart, beginning in March 2018, with all data collected by mid-July 2018 (within a 5-month period). Three mice from each treatment were included from each technical repeat (combined for 6 mice per treatment). No adverse events were experienced by mice in this sub-study. Four-week-old mice were fed a low-fat diet for four weeks (Figure 8). From eight weeks-of-age, mice in treatment 1 were fed a low-fat diet, while remaining mice were fed a high-fat diet (treatments 2-4). Mice were treated twice-a-week with: mock-irradiation (treatments 1 and 2) with vehicle; UVR then vehicle (treatment 3); UVR then cPTIO (UVR + cPTIO, treatment 4; cPTIO is a nitric oxide scavenger) (see Figure 8). Mice were fed diets and administered treatments for 12 weeks until 20 weeks-of-age.

**Figure 8.**
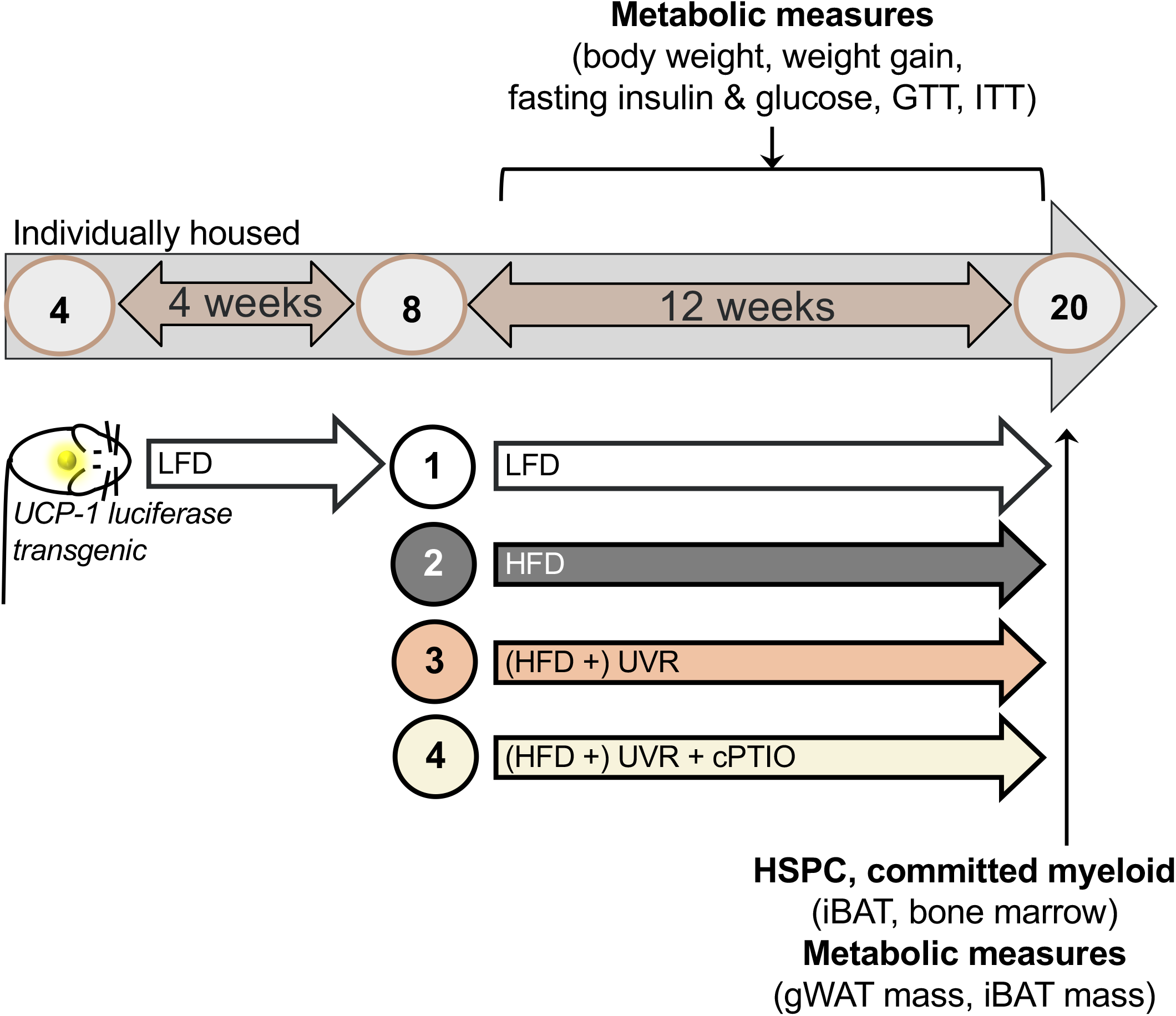
Experiment set-up. Four-week-old uncoupling-1 protein luciferase transgenic male mice were fed a low-fat diet (LFD) for four weeks. From eight weeks-of-age, mice were fed the LFD (treatment 1) or switched to the high-fat diet (HFD, treatments 2-4). From eight weeks-of-age, the shaved dorsal skin of mice was treated twice a week with: mock-irradiation and then vehicle applied to skin (treatments 1 and 2); sub-erythemal UVR (1 kJ m^-2^ UVB) and then vehicle (treatment 3); or, sub-erythemal UVR and then the nitric oxide scavenger cPTIO (0.1 mmol) was applied to skin (UVR + cPTIO, treatment 4). Mice were fed the LFD or HFD and administered the treatments twice a week for 12 weeks until 20 weeks-of-age. Here we report new findings of a sub-study of these previously published experiments^16^, in which we consider the relationships between metabolic dysfunction and hematopoietic stem and progenitor cell (HSPC) and committed myeloid cell phenotypes in interscapular brown adipose tissue (iBAT) and bone marrow for *n*=6 mice/treatment (*n*=24 in total). Two technical repeats of the same experiment were done with results combined, with three mice from each treatment per technical repeat combined for 6 mice per treatment. Mice were weighed weekly with body weights and weight gain calculated (after 12 weeks), with fasting insulin measured after 9 weeks, fasting glucose and glucose tolerance tests (GTT) performed after 10 weeks, and, insulin tolerance tests (ITT) performed after 11 weeks of dietary and skin treatments. Bone marrow, iBAT and gonadal white adipose tissue (gWAT) were dissected from mice at the end of the experiment, with HSPC and committed myeloid cells phenotyped in iBAT and bone marrow (via flow cytometry), and gWAT and iBAT weighed.

### Diet

All diets were obtained from Specialty Feeds (Glenn Forrest, Western Australia). Mice were fed either a high-fat diet (45.9% digestible energy as fat; SF12-031) or a low-fat diet (12.0% digestible energy as fat; SF12-029, control diet) from 4 weeks-of-age. Aside from changes in the macronutrient content, the low-fat diet was matched for nutrient content with the high-fat diet (see Supplementary table 5 to compare contents of both semi-purified diets).

### UVR

Mice with clean-shaven dorsal skin (8 cm^2^) were exposed (whole-body) to sub-erythemal/sub-oedemal UVB radiation (1 kJ m^-2^; 3.2 ± 0.3 min (mean ± SD)) emitted from lamps (held above dorsal surface of mice)^16, 23, 28, 35^. A bank of 40 W FS40 lamps (Philips TL UV-B, Eindhoven, The Netherlands) was used to administer UVR. These emit broadband UVR (250–360 nm), with 65% of the output in the UVB range (280–315 nm), and the remaining UVR in the UVC (250–280 nm) and UVA (315–360 nm) ranges^36^. Polyvinylchloride plastic was used to block wavelengths lower than 280 nm, as previously^16, 23, 28, 35^. Prior to each treatment, the time required to deliver the 1 kJ/m^2^ dose of UVB radiation was calculated using a handheld UV radiometer (UVX Digital Radiometer, Ultraviolet Products Inc., Upland, CA). Mice in the UVR treatments (groups 3 and 4, Figure 8) received a total dose of 24 kJ m^-2^ UVB across the 12 weeks of the experiment. The sub-erthymeal/sub-oedemal UVR dose (1 kJ m^-2^) was determined in initial experiments that tested skin responses to increasing doses of UVB (1, 2 or 4 kJ m^-2^), in which a micrometer (Mitutoyo Corp, Aurora, IL, USA) was used to measure back and ear skin swelling responses 24-48 h post-exposure. The 1 kJ m^-2^ dose (only) did not cause an oedemal response, with no skin swelling observed. Throughout the experiment, no adverse skin responses, including redness or irritation were observed in mice exposed to UVR. During each exposure session, mice were housed in individual compartments of a Perspex box. Mice were shaved at 8 weeks of age (prior to treatments commencing), and then every 2 weeks. For mock treatments, mice were handled in the same manner and held under standard, non-UVR emitting fluorescent lighting for the same time used to irradiate other mice.

### Topical skin treatment

The shaved dorsal skin some mice (treatment 4, Figure 8) was administered 0.1 mmol cPTIO (2-(4-Carboxyphenyl)-4,4,5,5-tetramethylimidazoline-1-oxyl-3-oxide potassium salt, Sigma-Aldrich (St Louis, MO, USA); a nitric oxide scavenger) immediately after delivery of UVR, in a vehicle (100 µL) as previously described^16, 22, 23^. This vehicle consisted of ethanol (Sigma-Aldrich), propylene glycol (Sigma-Aldrich) and double distilled water (2:1:1); 100 µL was administered to the shaved dorsal skin of mice in treatments 1-3 (Figure 8) immediately following exposure to UVR or mock treatment.

### Measuring weight gain and tissue weights

Mice were weighed weekly on Monday mornings, using a digital high accuracy bench micro-weighing scale (CAS MWP-3000H, >0.05 g sensitivity). The percentage weight gain was calculated from body weight measured at 8 weeks of age. At the conclusion of each experiment, mice were humanely euthanised via cervical dislocation following cardiac puncture for blood exsanguination (under isofluorane anaesthesia). An analytical scale (Analytical Standard Electric Balance, Ohaus, NJ, USA; >0.0001 g sensitivity) was used to weigh gWAT and iBAT dissected from mice at the end of the experiment.

### Glucose and insulin tolerance tests

Mice were transferred to a new cage (as singletons) and fasted for 5 h and then intraperitoneally challenged with either 1 g kg^-1^ glucose for glucose tolerance tests or with 0.75 IU kg^-1^ insulin for insulin tolerance tests (with no anaesthesia). Fasting glucose was measured before injection and at 15, 30, 45, 60, 90, and 120 min post-injection using Accu-Chek Performa glucometers and strips (Roche, Castle Hill, Australia) from the tail following initial removal of 1 cm end of the tail-tip.

### Flow cytometric analyses of HSPC and myeloid cells in bone marrow and iBAT

At the end of the experiment, cells from bone marrow and iBAT were phenotyped using flow cytometry as described previously^15^. All samples from each mouse were kept separate and not pooled. Bone marrow was isolated from long bones (femurs and tibias) and red blood cells lysed^15^. A single cell suspension of vascular stromal cells was isolated from minced iBAT (using scissors) digested for 90 min with collagenase IV (3 mg mL^-1^, Worthington, Lakewood NJ, USA) at 37 °C under gentle agitation^16, 37^. We have previously tested the activity of collagenase IV by ‘digesting’ bone marrow with this enzyme, observing no significant effects on extracellular marker expression by DC subsets and precursors^38^. In adipose tissue, both immune cells and HSPC reside in the vascular stromal cell fraction (non-adipose cells)^39^, including cells such as: adipose-derived stem/stromal cells, endothelial precursor cells, endothelial cells, macrophages, smooth muscle cells, lymphocytes, pericytes, and pre-adipocytes (among others)^40^. Vascular stromal iBAT cells and bone marrow cell numbers were determined after counting cells using a Neubauer chamber and excluding red blood cells and dead cells identified via staining with trypan blue. Previously optimised staining protocols^15^ were used to phenotype HSPC and myeloid cells using antibody panels detailed in the Supplementary file (Supplementary tables 6 and 7, respectively, including; target, fluorochrome, clone, supplier). Stem cell progenitors from the bone marrow of the genetic background (FVB/NJ) of mice used in the current study express Sca-1 and/or c-Kit at levels similar to C57Bl/6 mice^41^ (see also^12, 13^ for comparison to other similar published work for C57Bl/6 mice). Single cell suspensions for myeloid panel staining were incubated in Fc Block (Cat#553142, BD Biosciences, Franklin Lakes, NJ, USA). Fluorescent minus one staining controls were used for both panels where required. A 4-laser LSR Fortessa (BD Biosciences) flow cytometer was used to collect data, which were analysed using FlowJo software (v10.5.3 Mac OS X, BD Biosciences).

### Statistical analyses

All results were analyzed using GraphPad Prism (v8.4.3 for MAC OS10, 2020). Each mouse was considered an experimental unit. Descriptive statistics were calculated with mean and standard deviation (SD) reported for data in Figure 1g and Figure 1h, and mean and range for data in Figure 4-7. Area under the curve analyses of glucose tolerance test and insulin tolerance test data used zero as the baseline. All data were subjected to normality tests (D’Agostino & Pearson, Shapiro-Wilk) to determine if parametric data analyses were appropriate. ROUT (Q=1%) tests were performed to identify whether datapoints were significant outliers. Results were considered statistically significant for *P-values* < 0.05.

The proportions (or percentage) of each cell subtype within the total vascular stromal cell fraction of iBAT or total cells isolated from the long bones (bone marrow) were calculated via gating strategies (Supplementary figures 1,3-5). Cell proportion data considered changes to cell populations within the total isolated cell fraction, including stromal cells and red blood cells (independent of iBAT mass). For each cell subtype, we also compared the numbers of cells isolated from iBAT or bone marrow, which particularly for iBAT was done to consider differences induced by fat deposition (induced by high-fat diet), with cell numbers divided by the specific iBAT mass measured for each mouse. This was less relevant for bone marrow specimens, in which there is less ‘physical space’ for fat deposition^42^.

We first considered correlations within the whole dataset to understand the relationships between metabolic dysfunction and cell type. Data were combined as a whole dataset (*n*=24) and the linear relationships between metabolic and cell outcomes were evaluated using Pearson (if normally distributed) or Spearman (if not normally distributed; done for fasting insulin data only) correlation tests. We performed *n*=594 correlation tests, with 39 tests achieving *P-values* < 0.05. We did not correct for the multiple correlation tests performed to avoid excluding possible false negatives that could be worthy of investigation in future studies. We then analysed data between individual treatments to examine causality (diet, UVR, UVR + cPTIO (blocking UVR-induced nitric oxide)). For comparisons between treatments (*n*=6/treatment), one-way ANOVA with Tukey’s post-hoc (if normally distributed) or Kruskal-Wallis test with Dunn’s post-hoc (if not normally distributed) analyses were used to define differences between treatments, unless otherwise stated.

## Data availability

Data are available on request to authors.

## Acknowledgments

This research was supported by the Diabetes Research Foundation of Western Australia and the Telethon Kids Institute. SG was funded by an Al and Val Rosenstrauss Research Fellowship from the Rebecca L Cooper Foundation. RML was supported by a NHMRC Senior Research Fellowship. The study funders were not involved in the design of the study; the collection, analysis, and interpretation of data; writing the report; or the decision to submit the report for publication.Thank you to: Telethon Kids Institute Bioresources staff for day-to-day care of the mice; Dr Emily Barrick (Telethon Kids Institute) for animal welfare; Dr Paul Stevenson (Telethon Kids Institute) and Dr Emma De Jong (Telethon Kids Institute) for biostatistical advice around the need to correct for multiple testing.

## Author contributions

SG conceived and designed this study with input from KM, KP, MF, PH, RL, RW, VM and DS. KM and KP acquired and analysed the data for the study with help from SG and DS. KM performed all cell isolation and flow cytometric assays, including data collection and analyses. All authors have contributed towards the interpretation of findings from this study, have played a role in drafting the article or revising it critically for its intellectual content and have given their final approval for this version of the paper to be published. SG is the guarantor of this work.

