## Supplementary Information for "Metabolic dysfunction induced by high-fat diet modulates hematopoietic stem and myeloid progenitor cells in brown adipose tissue of mice"

**Supplementary table 1.** *Correlations between proportions of cell types in iBAT and metabolic outcomes*. The proportions of each cell subset were identified through the gating strategies of Supplementary figures 1 and 5. LSK=lineage$^-$Sca-1$^+$cKit$^+$ cells; LT-HSC=long-term hematopoietic stem cells; ST-HSC=short-term hematopoietic stem cells; MPP=multipotent progenitors; MP=myeloid progenitors; CMP=common myeloid progenitors; MEP=megakaryocyte-erythrocyte progenitors; GMP=granulocyte-macrophage progenitors; MDP=macrophage-dendritic cell progenitors; pre-pDC=pre-plasmacytoid dendritic cells; pre-cDC=pre-conventional dendritic cells; pDC=plasmacytoid dendritic cells; cDC=conventional dendritic cells; infDC=inflammatory dendritic cells; MDSC=myeloid-derived suppressor cells; NK cell=natural killer cells; GTT (AUC) (glucose tolerance test area under the curve); gWAT (gonadal white adipose tissue); iBAT (interscapular brown adipose tissue); ITT (AUC) (insulin tolerance test area under the curve).

| | **Metabolic outcomes** (r‡, *P-value*) | | | | | | | | |
|---|---|---|---|---|---|---|---|---|---|
| **Cell type** | Body weight | Weight gain | GTT (AUC) | gWAT mass | Liver mass | iBAT mass | Fasting insulin | Fasting glucose | ITT (AUC) |
| LSK | -0.12, 0.58 | -0.05, 0.81 | -0.12, 0.59 | -0.16, 0.47 | 0.15, 0.47 | -0.15, 0.48 | -0.06, 0.76 | 0.23, 0.28 | -0.03, 0.88 |
| LT-HSC | -0.11, 0.61 | -0.12, 0.57 | -0.19, 0.38 | -0.26, 0.21 | 0.16, 0.46 | -0.16, 0.45 | 0.01, 0.95 | 0.18, 0.41 | -0.04, 0.86 |
| ST-HSC | -0.07, 0.74 | 0.03, 0.88 | -0.04, 0.84 | 0.03, 0.90 | 0.14, 0.52 | -0.05, 0.81 | 0.07, 0.74 | 0.38, 0.07 | 0.02, 0.91 |
| MPP | -0.35, 0.10 | -0.16, 0.44 | 0.13, 0.53 | -0.26, 0.21 | 0.14, 0.51 | 0.02, 0.94 | -0.38, 0.07 | -0.13, 0.56 | -0.21, 0.34 |
| MP | -0.31, 0.14 | -0.20, 0.35 | -0.15, 0.50 | -0.39, 0.06 | 0.21, 0.33 | -0.28, 0.18 | -0.01, 0.95 | 0.09, 0.69 | -0.23, 0.29 |
| CMP | Infrequently detected (<0.0005% cells, not detected in 10/24 samples) | | | | | | | | |
| MEP | -0.31, 0.15 | -0.20, 0.34 | -0.19, 0.39 | -0.38, 0.06 | 0.22, 0.30 | -0.29, 0.17 | 0.05, 0.83 | 0.09, 0.68 | -0.25, 0.24 |
| GMP | Infrequently detected (<0.00005% cells, not detected in 20/24 samples) | | | | | | | | |
| MDP | Not detected in any sample | | | | | | | | |
| pre-pDC | Infrequently detected (<0.00005% cells, not detected in 10/24 samples) | | | | | | | | |
| pre-cDC | Infrequently detected (<0.0005% cells, not detected in 1/24 samples) | | | | | | | | |
| pDC | Infrequently detected (<0.00006% cells, not detected in 18/24 samples) | | | | | | | | |
| cDC | 0.41, ***0.049**** | 0.40, 0.051 | -0.13, 0.55 | 0.33, 0.11 | 0.11, 0.61 | 0.41, ***0.046**** | 0.01, 0.97 | -0.01, 0.94 | -0.13, 0.54 |
| infDC | 0.07, 0.74 | 0.38, 0.07 | -0.09, 0.68 | 0.29, 0.16 | 0.11, 0.60 | -0.13, 0.55 | -0.29, 0.17 | -0.15, 0.49 | -0.39, 0.06 |
| B cells | -0.23, 0.27 | 0.08, 0.71 | 0.21, 0.33 | -0.11, 0.60 | 0.05, 0.81 | -0.10, 0.65 | -0.32, 0.13 | -0.14, 0.50 | -0.24, 0.25 |
| MDSC | 0.12, 0.59 | -0.08, 0.73 | 0.17, 0.43 | -0.31, 0.14 | 0.14, 0.53 | 0.02, 0.91 | 0.25, 0.25 | 0.03, 0.88 | 0.06, 0.77 |
| NKp46+ | 0.05, 0.81 | 0.36, 0.08 | 0.21, 0.33 | 0.17, 0.42 | -0.08, 0.72 | -0.10, 0.65 | -0.31, 0.14 | -0.06, 0.79 | -0.11, 0.62 |
| CD3+ | -0.02, 0.91 | -0.01, 0.97 | 0.18, 0.41 | 0.01, 0.98 | -0.16, 0.47 | 0.18, 0.42 | -0.21, 0.34 | -0.06, 0.80 | 0.24, 0.28 |
| Neutrophils | -0.16, 0.46 | -0.14, 0.53 | -0.05, 0.82 | -0.42, ***0.041****† | 0.21, 0.32 | -0.20, 0.36 | 0.07, 0.75 | 0.04, 0.86 | -0.30, 0.16 |

***P-values < 0.05*** with Pearson correlation tests were performed for all comparisons, except for fasting insulin as normality tests indicated that these data were not normally distributed, for which the Spearman correlation test was used instead. †One datapoint (within the percentages of neutrophils in iBAT dataset) was identified as an outlier (via ROUT test), and after the exclusion of this datapoint, this correlation become non-significant.

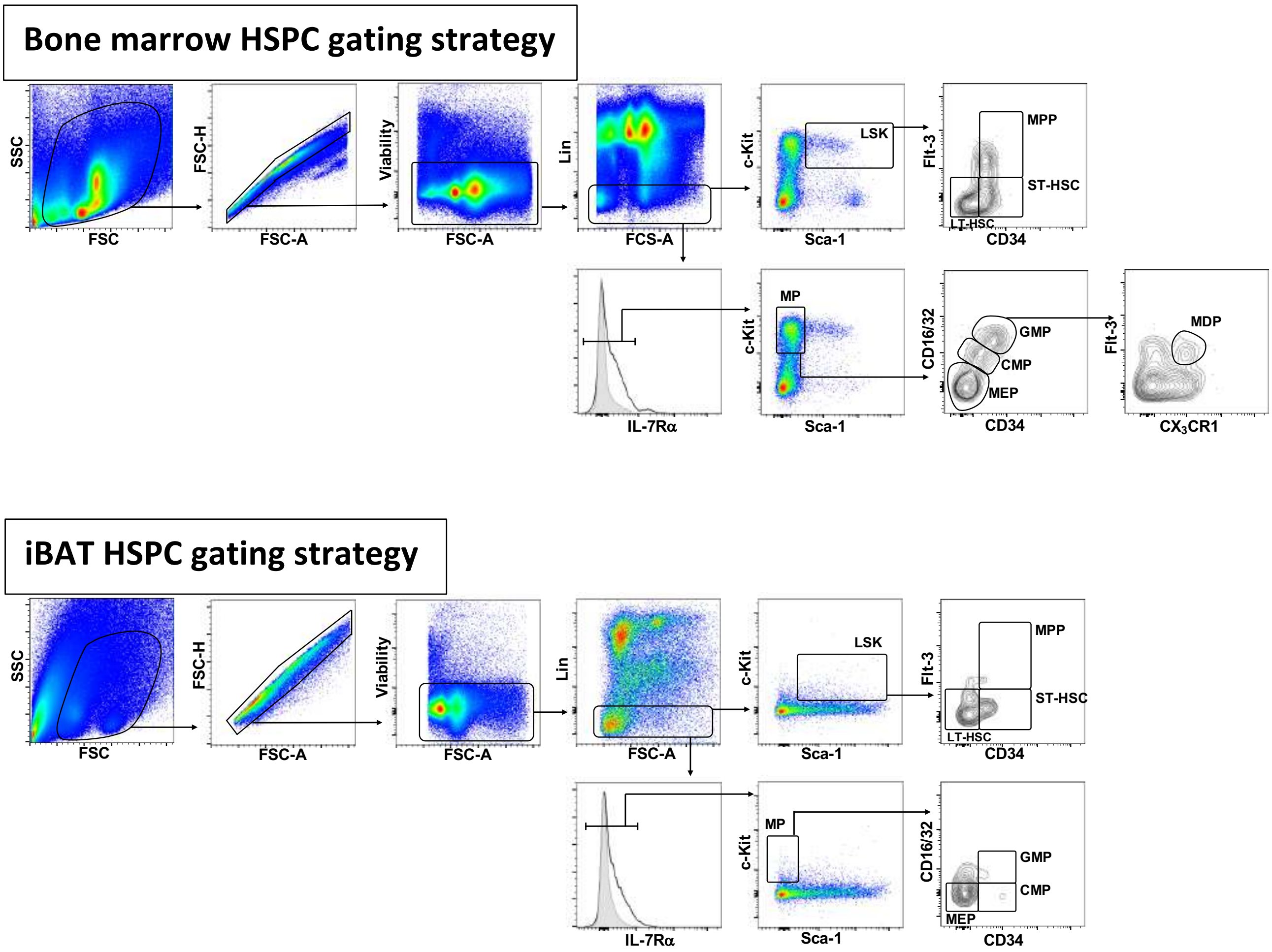


**Supplementary figure 1:** Phenotyping FlowJo® gating strategy used to select for HSPC in bone marrow and iBAT via multi-colour flow cytometry. GMP=granulocyte-macrophage progenitors; CMP=common myeloid progenitors; MEP=megakaryocyte-erythrocyte progenitors; MDP=macrophage-dendritic cell progenitors; MP=myeloid progenitors; LSK=lineage$^-$Sca-1$^+$cKit$^+$ cells; MPP=multipotent progenitors; ST-HSC=short-term hematopoietic stem cells. Shaded area in histograms represents the IL-7Ra FMO (fluorescence minus one) control.

**Supplementary table 2.** *Correlations between numbers of cell types in iBAT (per g of tissue) and metabolic outcomes.* The numbers of each cell subset were calculated by multiplying the total number of live vascular stromal cells (excluding red blood and dead cells), with proportions of specific cell types identified through gating strategies of Supplementary figures 1 and 5. LSK=lineage$^{-}$Sca-1$^{+}$cKit$^{+}$ cells; LT-HSC=long-term hematopoietic stem cells; ST-HSC=short-term hematopoietic stem cells; MPP=multipotent progenitors; MP=myeloid progenitors; CMP=common myeloid progenitors; MEP=megakaryocyte-erythrocyte progenitors; GMP=granulocyte-macrophage progenitors; MDP=macrophage-dendritic cell progenitors; pre-pDC=pre-plasmacytoid dendritic cells; pre-cDC=pre-conventional dendritic cells; pDC=plasmacytoid dendritic cells; cDC=conventional dendritic cells; infDC=inflammatory dendritic cells; MDSC=myeloid-derived suppressor cells; NK cell=natural killer cells; GTT (AUC) (glucose tolerance test area under the curve); gWAT (gonadal white adipose tissue); iBAT (interscapular brown adipose tissue); ITT (AUC) (insulin tolerance test area under the curve).

| | **Metabolic outcomes** (r‡, *P-value*) | | | | | | | | |
|---|---|---|---|---|---|---|---|---|---|
| **Cell type** | Body weight | Weight gain | GTT (AUC) | gWAT mass | Liver mass | iBAT mass | Fasting insulin | Fasting glucose | ITT (AUC) |
| Total | -0.30, 0.16 | -0.22, 0.30 | 0.07, 0.74 | -0.37, 0.08 | -0.12, 0.56 | -0.39, 0.06 | -0.05, 0.83 | 0.03, 0.89 | 0.02, 0.92 |
| LSK | -0.18, 0.39 | -0.11, 0.62 | -0.08, 0.73 | -0.27, 0.21 | 0.12, 0.57 | -0.38, 0.06 | -0.06, 0.80 | 0.17, 0.41 | -0.09, 0.67 |
| LT-HSC | -0.20, 0.34 | -0.18, 0.39 | -0.13, 0.53 | -0.38, 0.07 | 0.14, 0.63 | -0.41, ***0.045**** | 0.01, 0.95 | 0.16, 0.47 | -0.11, 0.62 |
| ST-HSC | -0.10, 0.64 | -0.00, 0.99 | -0.03, 0.90 | -0.06, 0.79 | 0.15, 0.47 | -0.23, 0.27 | 0.02, 0.92 | 0.32, 0.13 | -0.03, 0.91 |
| MPP | -0.31, 0.14 | -0.14, 0.51 | 0.22, 0.29 | -0.21, 0.31 | 0.03, 0.90 | -0.08, 0.73 | -0.41, ***0.047**** | -0.16, 0.47 | -0.14, 0.52 |
| MP | -0.32, 0.13 | -0.24, 0.27 | -0.08, 0.72 | -0.46, ***0.02**** | 0.08, 0.70 | -0.45, ***0.03**** | -0.04, 0.87 | 0.05, 0.81 | -0.17, 0.42 |
| CMP | Infrequently detected (<20 cells/g tissue, not detected in 10/24 samples) | | | | | | | | |
| MEP | -0.33, 0.12 | -0.25, 0.24 | -0.13, 0.56 | -0.48, ***0.02****† | 0.10, 0.63 | -0.50, ***0.01**** | -0.02, 0.91 | 0.05, 0.83 | -0.21, 0.33 |
| GMP | Infrequently detected (<5 cells/g tissue, not detected in 20/24 samples) | | | | | | | | |
| MDP | Not detected in any sample | | | | | | | | |
| pre-pDC | Infrequently detected (<30 cells/g tissue, not detected in 10/24 samples) | | | | | | | | |
| pre-cDC | Infrequently detected (<100 cells/g tissue, not detected in 1/24 samples) | | | | | | | | |
| pDC | Infrequently detected (<10 cells/g tissue, not detected in 18/24 samples) | | | | | | | | |
| cDC | 0.26, 0.23 | 0.31, 0.14 | 0.07, 0.74 | 0.20, 0.36 | -0.04, 0.86 | 0.14, 0.50 | -0.06, 0.77 | -0.04, 0.84 | 0.04, 0.86 |
| infDC | -0.05, 0.83 | 0.25, 0.23 | 0.02, 0.93 | 0.11, 0.62 | 0.03, 0.90 | -0.29, 0.17 | -0.23, 0.28 | -0.24, 0.27 | -0.27, 0.21 |
| B cell | -0.28, 0.18 | -0.08, 0.72 | 0.22, 0.30 | -0.26, 0.22 | -0.15, 0.50 | -0.21, 0.32 | -0.30, 0.15 | -0.13, 0.55 | -0.04, 0.87 |
| MDSC | -0.18, 0.41 | -0.30, 0.16 | 0.16, 0.47 | -0.54, ***0.006**** | -0.00, 1.00 | -0.21, 0.33 | 0.16, 0.46 | 0.07, 0.73 | 0.03, 0.89 |
| NKp46+ | -0.05, 0.80 | 0.16, 0.45 | 0.21, 0.31 | -0.05, 0.80 | -0.14, 0.51 | -0.22, 0.31 | -0.29, 0.17 | -0.10, 0.66 | 0.03, 0.89 |
| CD3+ | -0.10, 0.64 | -0.09, 0.67 | 0.20, 0.35 | -0.16, 0.45 | -0.05, 0.82 | 0.01, 0.98 | -0.23, 0.28 | 0.03, 0.89 | 0.08, 0.72 |
| Neutrophils | -0.32, 0.13 | -0.34, 0.10 | 0.03, 0.91 | -0.59, ***0.002****†† | 0.06, 0.78 | -0.31, 0.13 | 0.01, 0.95 | 0.07, 0.74 | -0.16, 0.46 |

***P-values < 0.05*** with Pearson correlation tests were performed for all comparisons, except for fasting insulin as normality tests indicated that these data were not normally distributed, for which the Spearman correlation test was used instead. †While no outliers were identified (via ROUT test), the exclusion of one 'more extreme' datapoint from the MEP dataset (identified in Figure 2g) changed the outcome of this correlation to non-significant. ††Two datapoints (within the numbers of neutrophils in iBAT dataset) were identified as outliers (via ROUT test), and after their exclusion this correlation became non-significant.

**Supplementary table 3.** *Correlations between proportions of cell types in bone marrow and metabolic outcomes*. The proportions of each cell subset were identified through gating strategies of Supplementary figures 1 and 4. LSK=lineage$^-$Sca-1$^+$cKit$^+$ cells; LT-HSC=long-term hematopoietic stem cells; ST-HSC=short-term hematopoietic stem cells; MPP=multipotent progenitors; MP=myeloid progenitors; CMP=common myeloid progenitors; MEP=megakaryocyte-erythrocyte progenitors; GMP=granulocyte-macrophage progenitors; MDP=macrophage-dendritic cell progenitors; pre-pDC=pre-plasmacytoid dendritic cells; pre-cDC=pre-conventional dendritic cells; pDC=plasmacytoid dendritic cells; cDC=conventional dendritic cells; infDC=inflammatory dendritic cells; MDSC=myeloid-derived suppressor cells; NK cell=natural killer cells; GTT (AUC) (glucose tolerance test area under the curve); gWAT (gonadal white adipose tissue); iBAT (interscapular brown adipose tissue); ITT (AUC) (insulin tolerance test area under the curve).

| | **Metabolic outcomes** (r[‡], *P-value*) | | | | | | | | |
|---|---|---|---|---|---|---|---|---|---|
| **Cell type** | Body weight | Weight gain | GTT (AUC) | gWAT mass | Liver mass | iBAT mass | Fasting insulin | Fasting glucose | ITT (AUC) |
| LSK | 0.03, 0.87 | 0.15, 0.47 | -0.31, 0.13 | 0.24, 0.26 | 0.14, 0.51 | -0.10, 0.64 | -0.32, 0.12 | -0.34, 0.10 | -0.21, 0.33 |
| LT-HSC | -0.03, 0.90 | 0.05, 0.80 | -0.29, 0.16 | 0.17, 0.42 | 0.11, 0.61 | 0.08, 0.72 | -0.44, **_0.03*_** | -0.38, 0.07 | -0.22, 0.30 |
| ST-HSC | -0.04, 0.86 | 0.02, 0.94 | -0.25, 0.24 | 0.10, 0.66 | 0.32, 0.12 | -0.42, **_0.04*_** | -0.31, 0.14 | -0.16, 0.46 | -0.20, 0.35 |
| MPP | -0.18, 0.40 | 0.05, 0.80 | 0.17, 0.42 | 0.13, 0.53 | -0.20, 0.34 | -0.07, 0.74 | -0.64, **_0.0008*_** | -0.12, 0.58 | -0.22, 0.31 |
| MP | 0.20, 0.36 | 0.06, 0.79 | 0.04, 0.85 | 0.13, 0.54 | 0.12, 0.59 | -0.07, 0.76 | -0.17, 0.43 | -0.40, 0.05 | 0.18, 0.39 |
| CMP | 0.10, 0.66 | 0.07, 0.75 | 0.25, 0.24 | 0.23, 0.28 | 0.12, 0.59 | -0.16, 0.46 | -0.15, 0.48 | -0.17, 0.43 | 0.07, 0.74 |
| MEP | 0.30, 0.15 | 0.08, 0.73 | 0.01, 0.95 | 0.06, 0.79 | 0.05, 0.82 | 0.12, 0.56 | -0.17, 0.44 | -0.52, **_0.009*_** | 0.29, 0.18 |
| GMP | -0.08, 0.70 | -0.04, 0.87 | 0.07, 0.74 | 0.15, 0.48 | 0.19, 0.36 | -0.35, 0.09 | -0.21, 0.33 | -0.00, 0.98 | -0.05, 0.83 |
| MDP | -0.23, 0.29 | -0.04, 0.86 | 0.22, 0.30 | 0.10, 0.63 | -0.26, 0.22 | -0.05, 0.82 | -0.66, **_0.0005*_** | -0.17, 0.43 | -0.13, 0.53 |
| pre-pDC | 0.13, 0.53 | -0.12, 0.59 | 0.29, 0.17 | -0.22, 0.31 | -0.00, 0.99 | 0.22, 0.31 | 0.17, 0.41 | 0.20, 0.34 | 0.39, 0.06 |
| pre-cDC | -0.07, 0.75 | -0.38, 0.07 | 0.23, 0.27 | -0.40, 0.05 | 0.32, 0.13 | -0.00, 0.99 | 0.23, 0.27 | 0.08, 0.70 | 0.22, 0.30 |
| pDC | -0.06, 0.78 | 0.06, 0.80 | -0.12, 0.56 | 0.05, 0.80 | 0.16, 0.46 | 0.08, 0.73 | -0.07, 0.76 | 0.11, 0.60 | -0.34, 0.10 |
| cDC | 0.15, 0.49 | 0.16, 0.46 | 0.01, 0.97 | 0.02, 0.94 | -0.06, 0.76 | 0.13, 0.53 | 0.59, **_0.003*_** | 0.50, **_0.01*_** | 0.07, 0.73 |
| infDC | 0.43, **_0.04*_** | 0.54, **_0.006*_** | -0.08, 0.71 | 0.50, **_0.01*_** | -0.03, 0.88 | 0.25, 0.24 | 0.20, 0.34 | 0.11, 0.59 | -0.00, 1.00 |
| B cell | 0.07, 0.74 | 0.24, 0.25 | 0.04, 0.86 | 0.15, 0.49 | 0.15, 0.49 | -0.18, 0.40 | 0.27, 0.19 | 0.25, 0.24 | -0.29, 0.18 |
| MDSC | -0.33, 0.12 | -0.48, **_0.02*†_** | 0.02, 0.94 | -0.50, **_0.01*†_** | 0.04, 0.84 | -0.25, 0.24 | 0.37, 0.07 | 0.40, 0.05 | 0.09, 0.67 |
| NKp46+ | -0.21, 0.32 | -0.15, 0.50 | 0.27, 0.20 | -0.01, 0.97 | -0.30, 0.15 | 0.15, 0.48 | -0.37, 0.07 | -0.05, 0.81 | 0.18, 0.39 |
| CD3+ | 0.19, 0.37 | 0.10, 0.63 | 0.16, 0.47 | -0.03, 0.89 | 0.22, 0.30 | 0.05, 0.81 | 0.10, 0.63 | 0.03, 0.87 | 0.13, 0.53 |

**_*P-values < 0.05_** with Pearson correlation tests were performed for all comparisons, except for fasting insulin as normality tests indicated that these data were not normally distributed, for which the Spearman correlation test was used instead. [†]While no outliers were identified (via ROUT test), the exclusion of one 'more extreme' datapoint (identified in Figure 3j) changed the outcome of some correlations to non-significant (denoted via[†]).

**Supplementary table 4.** *Correlations between numbers of cell types in bone marrow and metabolic outcomes.* The numbers of each cell subset were calculated by multiplying the total number of live bone marrow cells (excluding red blood and dead cells), with proportions of specific cell types identified through gating strategies of Supplementary figures 1 and 4. LSK=lineage$^-$Sca-1$^+$cKit$^+$ cells; LT-HSC=long-term hematopoietic stem cells; ST-HSC=short-term hematopoietic stem cells; MPP=multipotent progenitors; MP=myeloid progenitors; CMP=common myeloid progenitors; MEP=megakaryocyte-erythrocyte progenitors; GMP=granulocyte-macrophage progenitors; MDP=macrophage-dendritic cell progenitors; pre-pDC=pre-plasmacytoid dendritic cells; pre-cDC=pre-conventional dendritic cells; pDC=plasmacytoid dendritic cells; cDC=conventional dendritic cells; infDC=inflammatory dendritic cells; MDSC=myeloid-derived suppressor cells; NK cell=natural killer cells; GTT (AUC) (glucose tolerance test area under the curve); gWAT (gonadal white adipose tissue); iBAT (interscapular brown adipose tissue); ITT (AUC) (insulin tolerance test area under the curve).

| | **Metabolic outcomes** (r[‡], *P-value*) | | | | | | | | |
|---|---|---|---|---|---|---|---|---|---|
| **Cell type** | Body weight | Weight gain | GTT (AUC) | gWAT mass | Liver mass | iBAT mass | Fasting insulin | Fasting glucose | ITT (AUC) |
| Total | 0.02, 0.91 | 0.01, 0.97 | -0.24, 0.26 | 0.21, 0.32 | -0.06, 0.78 | 0.22, 0.30 | -0.07, 0.73 | -0.27, 0.19 | -0.19, 0.39 |
| LSK | -0.25, 0.24 | -0.08, 0.71 | -0.10, 0.65 | -0.06, 0.80 | 0.07, 0.76 | -0.17, 0.43 | -0.31, 0.14 | -0.43, ***0.03**** † | -0.32, 0.13 |
| LT-HSC | -0.00, 0.99 | 0.05, 0.81 | -0.37, 0.08 | 0.19, 0.37 | 0.12, 0.60 | 0.11, 0.61 | -0.28, 0.20 | -0.38, 0.07 | -0.26, 0.23 |
| ST-HSC | -0.30, 0.16 | -0.17, 0.43 | -0.14, 0.51 | -0.09, 0.68 | 0.23, 0.27 | -0.35, 0.09 | -0.31, 0.14 | -0.36, 0.09 | -0.37, 0.07 |
| MPP | -0.33, 0.12 | -0.08, 0.72 | 0.11, 0.61 | -0.04, 0.87 | -0.12, 0.58 | -0.18, 0.41 | -0.55, ***0.005**** | -0.32, 0.13 | -0.32, 0.13 |
| MP | -0.13, 0.54 | -0.12, 0.57 | -0.05, 0.83 | -0.05, 0.83 | 0.06, 0.79 | -0.09, 0.69 | -0.36, 0.09 | -0.62, ***0.001**** | -0.16, 0.45 |
| CMP | -0.24, 0.26 | -0.13, 0.54 | 0.08, 0.73 | -0.00, 1.00 | 0.03, 0.87 | -0.17, 0.42 | -0.34, 0.10 | -0.45, ***0.03**** † | -0.25, 0.23 |
| MEP | 0.06, 0.78 | -0.05, 0.80 | -0.09, 0.67 | -0.05, 0.82 | 0.02, 0.91 | 0.11, 0.62 | -0.32, 0.12 | -0.72, ***<0.0001**** | -0.00, 1.00 |
| GMP | -0.35, 0.09 | -0.19, 0.36 | 0.03, 0.90 | -0.05, 0.84 | 0.11, 0.61 | -0.33, 0.12 | -0.33, 0.11 | -0.30, 0.16 | -0.31, 0.14 |
| MDP | -0.34, 0.10 | -0.12, 0.56 | 0.12, 0.58 | -0.07, 0.74 | -0.12, 0.57 | -0.17, 0.43 | -0.55, ***0.005**** | -0.34, 0.10 | -0.28, 0.19 |
| pre-pDC | 0.07, 0.74 | -0.18, 0.40 | 0.21, 0.33 | -0.24, 0.26 | -0.08, 0.73 | 0.27, 0.21 | 0.10, 0.66 | 0.05, 0.81 | 0.35, 0.09 |
| pre-cDC | -0.25, 0.24 | -0.45, ***0.03**** † | 0.13, 0.54 | -0.40, 0.06 | 0.31, 0.14 | -0.09, 0.68 | 0.13, 0.54 | -0.02, 0.92 | 0.02, 0.94 |
| pDC | -0.29, 0.17 | -0.12, 0.59 | -0.08, 0.70 | -0.09, 0.67 | 0.09, 0.68 | -0.04, 0.84 | -0.09, 0.68 | -0.28, 0.19 | -0.45, ***0.03**** |
| cDC | 0.03, 0.91 | 0.09, 0.68 | -0.05, 0.81 | 0.04, 0.87 | -0.04, 0.85 | 0.14, 0.50 | 0.45, ***0.03**** | 0.18, 0.39 | -0.14, 0.52 |
| infDC | -0.01, 0.95 | 0.05, 0.48 | -0.01, 0.95 | 0.11, 0.60 | -0.03, 0.88 | 0.01, 0.95 | 0.11, 0.60 | -0.25, 0.23 | -0.23, 0.28 |
| B cell | 0.05, 0.82 | 0.25, 0.26 | -0.05, 0.82 | 0.31, 0.15 | 0.03, 0.89 | -0.08, 0.72 | 0.33, 0.13 | 0.22, 0.30 | -0.38, 0.08 |
| MDSC | -0.38, 0.07 | -0.28, 0.19 | 0.01, 0.95 | -0.23, 0.29 | 0.00, 0.99 | -0.15, 0.47 | 0.09, 0.68 | -0.25, 0.23 | -0.25, 0.24 |
| NKp46+ | -0.36, 0.08 | -0.17, 0.42 | 0.15, 0.48 | -0.10, 0.65 | -0.19, 0.39 | -0.09, 0.69 | -0.44, ***0.03**** † | -0.31, 0.14 | -0.15, 0.47 |
| CD3+ | 0.23, 0.28 | 0.19, 0.39 | -0.11, 0.61 | 0.28, 0.20 | 0.12, 0.60 | 0.21, 0.33 | 0.32, 0.14 | -0.08, 0.72 | -0.10, 0.66 |

***P-values < 0.05*** with Pearson correlation tests were performed for all comparisons, except for fasting insulin as normality tests indicated that these data were not normally distributed, for which the Spearman correlation test was used instead. [†]While no outliers were identified (via ROUT test), the exclusion of one 'more extreme' datapoint (identified in Supplementary figure 2) changed the outcome of some correlations to non-significant (denoted via[†]).

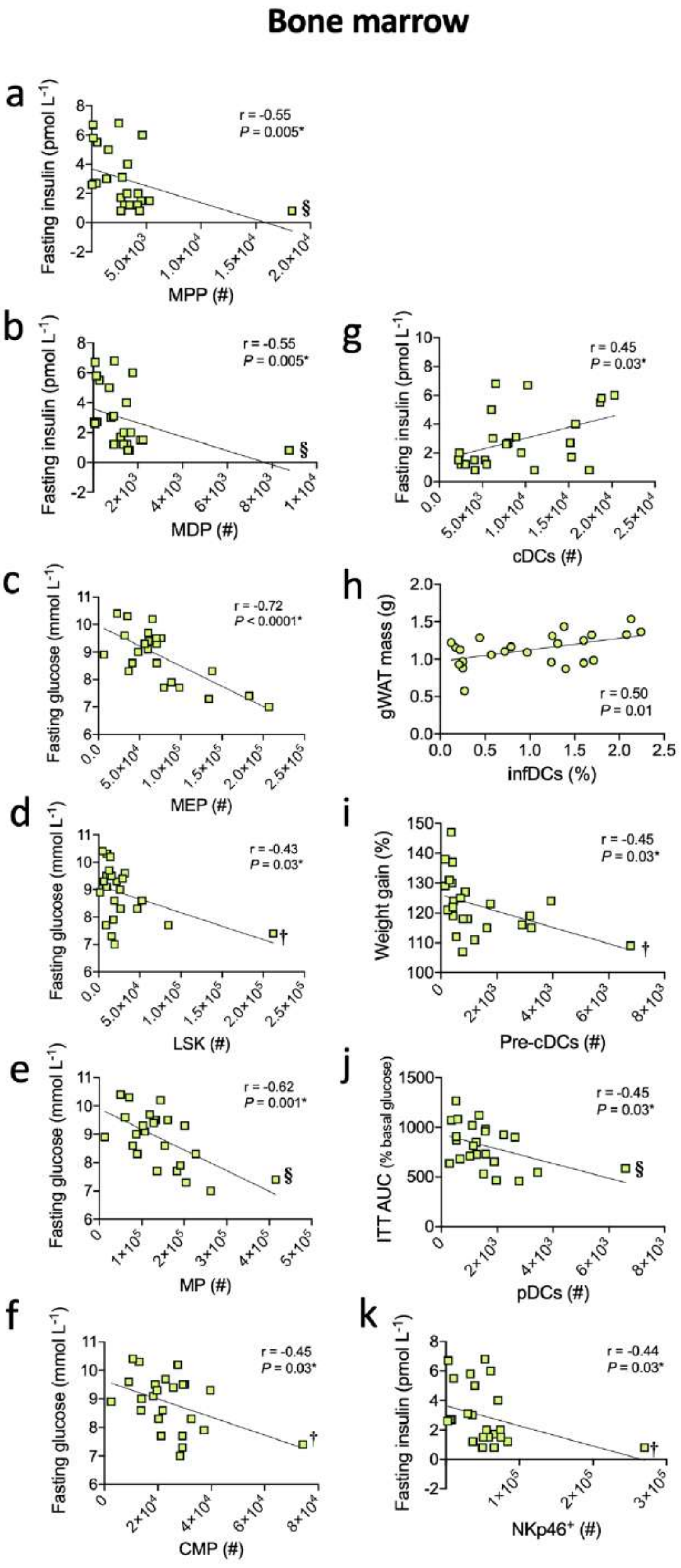


**Supplementary figure 2:** *Increasing metabolic dysfunction and adiposity is associated with declines in the numbers of some HSPC subsets in bone marrow.* Inverse linear relationships were observed between fasting insulin and the numbers of:

(**a**) multipotent progenitors (MPP), or
(**b**) macrophage-dendritic cell progenitor (MDP) in bone marrow.

Inverse linear relationships were observed between fasting glucose and numbers of:

(**c**) megakaryocyte-erythrocyte progenitors (MEP),
(**d**) lineage$^{-}$Sca-1$^{+}$cKit$^{+}$ cells (LSK),
(**e**) myeloid progenitors (MP), or
(**f**) common myeloid progenitors (CMP).

Positive linear relationships were observed between:

(**g**) numbers of conventional DC (cDC) and fasting insulin; and,
(**h**) proportions of inflammatory DC (infDC) and gonadal white adipose tissue (gWAT) mass.

Other inverse linear relationships were observed between numbers of:

(**i**) pre-conventional dendritic cells (pre-cDC) and weight gain;
(**j**) plasmacytoid DCs (pDCs) and insulin tolerance test area under the curve (ITT AUC) values; and,
(**k**) NKp46+ cells and fasting insulin.

Data were combined from 4 treatment groups of the experiment described in Figure 8 from *n*=24 individual mice. Two technical repeats of the same experiment were done with results combined, with three mice from each treatment per technical repeat combined for 6 mice per treatment. All data were compared using Pearson correlation (r) test (*p<0.05), except for fasting insulin data, which were compared using a Spearman correlation test as these data were not normally distributed. No outliers were identified (via ROUT test), although the exclusion of a 'more extreme' datapoint (†) in (d), (f), (i) or (k) changed the outcome of these correlations to non-significant [(r = -0.26, *P* = 0.09), (r = -0.34, *P* = 0.11), (r = -0.36, *P* = 0.09), or (r = -0.37, *P* = 0.08), respectively]. Conversely, following the exclusion of a similar datapoint (§) from (a), (b), (e) or (j), the correlations remained significant [(r = -0.50, *P* = 0.01), (r = -0.50, *P* = 0.02), (r = -0.59, *P* = 0.003), or (r =-0.46, *P* = 0.03), respectively].

**Supplementary table 5:** Nutritional composition of diets. This table contains the formulations for the high-fat and low-fat semi-purified diets.

| **Ingredients** (g $100g^{-1}$) | **High-Fat Diet** (SF12-031) | **Low-Fat Diet** (SF12-029) |
|---|---|---|
| Sucrose | 10.0 | 10.0 |
| Casein (acid) | 20.0 | 20.0 |
| Canola oil | 2.9 | 5.0 |
| Lard† | 20.7 | 0.0 |
| Cellulose | 5.0 | 5.0 |
| Wheat starch | 17.4 | 36.0 |
| Dextrinised starch | 13.2 | 13.2 |
| DL-methionine | 0.3 | 0.3 |
| AIN93_trace minerals | 0.14 | 0.14 |
| Lime (calcium carbonate) | 2.5 | 2.5 |
| Salt (fine sodium chloride) | 0.26 | 0.26 |
| Potassium dihydrogen phosphate | 0.76 | 0.76 |
| Potassium sulphate | 0.16 | 0.16 |
| Potassium citrate | 0.15 | 0.15 |
| Magnesium oxide | 0.17 | 0.17 |
| Dicalcium phosphate | 5.1 | 5.1 |
| AIN93 Vitamins (No vitamin D)‡ | 1.0 | 1.0 |
| Choline chloride 75% w/w | 0.25 | 0.25 |
| Food colour | 0.02 | 0.02 |
| % Digestible energy - lipid | 45.9% | 12.0% |
| % Digestible energy - protein | 17.7% | 22.0% |
| Digestible energy | 19 MJ $kg^{-1}$ | 15 MJ $kg^{-1}$ |

†As previously[1], the lard fraction of the high-fat diet likely contains vitamin D; however, the precise amount is unknown.

‡American Institute of Nutrition (AIN) trace minerals number 93 is a nutritional supplement used in rodent food designed to promote maintenance of health in adult rodents[2], to which vitamin D was not included.

**Supplementary table 6:** Hematopoietic stem and progenitor cell multi-colour flow cytometry panel

| Antibody | Fluorochrome | Clone | Supplier |
|---|---|---|---|
| Rat Anti-Mouse CD2 | Biotin | RM2-5 | BD Biosciences[1] |
| Hamster Anti-Mouse CD3 | Biotin | 145-2C11 | BD Biosciences |
| Rat Anti-Mouse CD4 | Biotin | GKl.5 | BD Biosciences |
| Rat Anti-Mouse CD5 | Biotin | 53-7.3 | BD Biosciences |
| Rat Anti-Mouse CD8α | Biotin | 53-6.7 | BD Biosciences |
| Rat Anti-Mouse CD19 | Biotin | 1D3 | BD Biosciences |
| Rat Anti-Mouse B220 (CD45R) | Biotin | RA3-6B2 | BD Biosciences |
| Rat Anti-Mouse Gr-1 | Biotin | RB6-8CS | BD Biosciences |
| Rat Anti-Mouse Ter119 | Biotin | TER-119 | BD Biosciences |
| Rat Anti-Mouse CD16/32 | PerCP-Cy5.5 | 2.4G2 | BD Biosciences |
| Rat Anti-Mouse CD34 | FITC | RAM34 | BD Biosciences |
| Rat Anti-Mouse IL-7Rα (CD127) | PE-Cy7 | SB/199 | BD Biosciences |
| Rat Anti-Mouse Flt-3 (CD135) | PE | A2F10.1 | BD Biosciences |
| Rat Anti-Mouse c-Kit (CD117) | APC-Cy7 | 2B8 | BD Biosciences |
| Rat Anti-Mouse Sca-1 | BV510 | D7 | BD Biosciences |
| Rat Anti-Mouse CX3CR1 | APC | SA011F11 | BioLegend[2] |
| Rat Anti-Mouse NKG2D | BV711 | CX5 | BD Biosciences |
| Viability | AF700 | - | BD Biosciences |
| Streptavidin | BV605 | - | BD Biosciences |

[1]BD Biosciences (Franklin Lakes, NJ, USA)
[2]Biolegend (San Diego, CA, USA)

**Supplementary table 7:** Myeloid cell multicolour flow cytometry panel

| Antibody | Fluorochrome | Clone | Supplier |
|---|---|---|---|
| Rat Anti-Mouse CD3 | FITC | 17A2 | BD Biosciences |
| Rat Anti-Mouse CD11b | BV510 | M1/70 | BD Biosciences |
| Hamster Anti-Mouse CD11c | BV711 | HL3 | BD Biosciences |
| Rat Anti-Mouse CD19 | APC-H7 | 1D3 | BD Biosciences |
| Rat Anti-Mouse CD24 | PE | M1/69 | BD Biosciences |
| Rat Anti-Mouse Gr-1 | Biotin | RB6-6B2 | BD Biosciences |
| Rat Anti-Mouse B220 (CD45R) | PerCP-Cy5.5 | RA3-6B2 | BD Biosciences |
| Rat Anti-Mouse NKp46 (CD335) | PE-Cy7 | 29A1.4 | BioLegend |
| Rat Anti-Mouse SIRP-α (CD172α) | APC | P84 | BioLegend |
| Rat Anti-Mouse I-A/I-E | BV421 | M5/114.15.2 | BioLegend |
| Rat Anti-Mouse F4/80 | BV785 | BM8 | BioLegend |
| Viability | AF700 | - | BD Biosciences |
| Streptavidin | BV605 | - | BD Biosciences |

[1]BD Biosciences (Franklin Lakes, NJ, USA)
[2]Biolegend (San Diego, CA, USA)

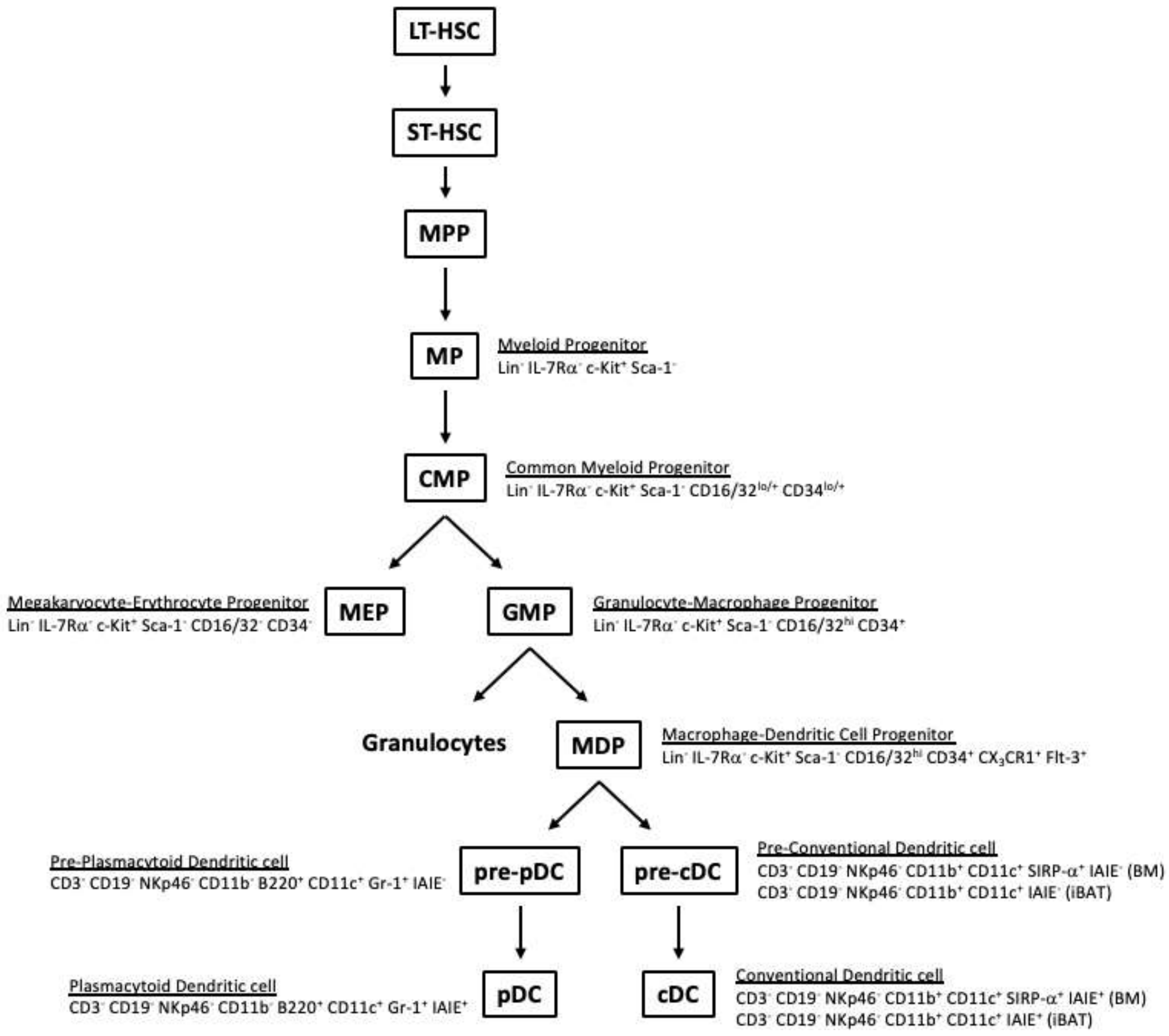


**Supplementary figure 3:** Hierarchical commitment of HSPCs to pDCs and cDCs. Shown is the developmental pathway within bone marrow resulting in the generation of terminally committed granulocytes, pDCs and cDCs and the associated cellular markers used to identify individual cell types. LT-HSC=long-term hematopoietic stem cells; ST-HSC=short-term hematopoietic stem cells; MPP=multipotent progenitors; MP=myeloid progenitors; CMP=common myeloid progenitors; MEP=megakaryocyte-erythrocyte progenitors; GMP=granulocyte-macrophage progenitors; MDP=macrophage-dendritic cell progenitors; pre-pDC=pre-plasmacytoid dendritic cells; pre-cDC=pre-conventional dendritic cells; pDC=plasmacytoid dendritic cells; cDC=conventional dendritic cells.

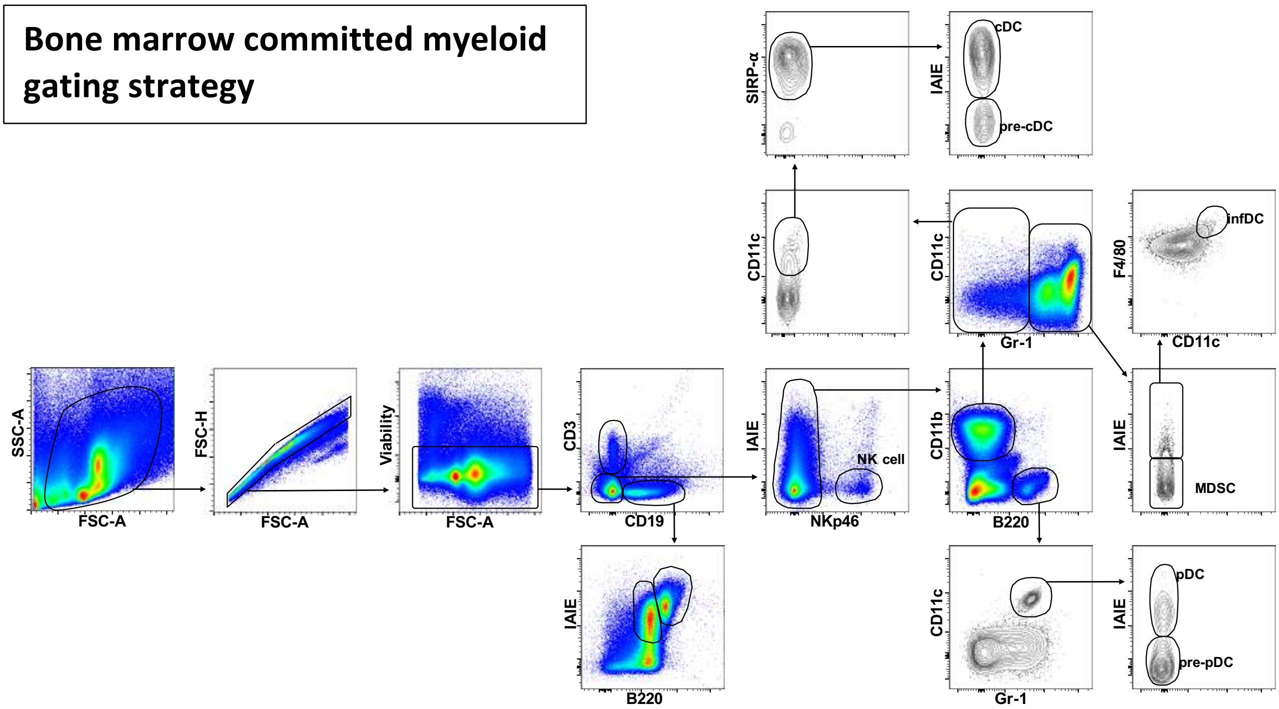


**Supplementary figure 4:** Phenotyping FlowJo® gating strategy used to select for committed myeloid cells in bone marrow via multi-colour flow cytometry. cDC=conventional dendritic cells; pre-cDC=pre-conventional dendritic cells; infDC=inflammatory dendritic cells; NK cell=natural killer cells; MDSC=myeloid-derived suppressor cells; pDC=plasmacytoid dendritic cells; pre-pDC=pre-plasmacytoid dendritic cells.

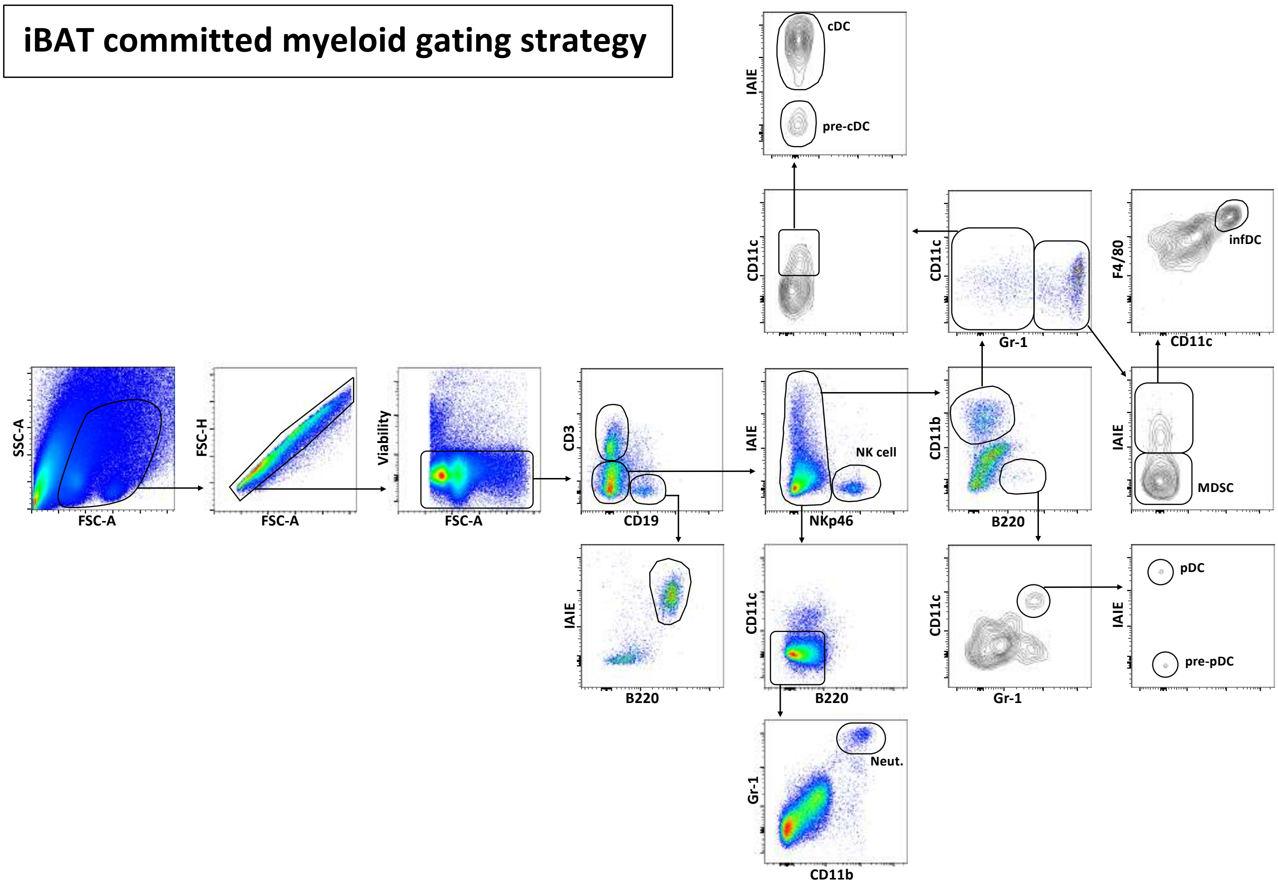


**Supplementary figure 5:** Phenotyping FlowJo® gating strategy used to select for committed myeloid cells in iBAT via multi-colour flow cytometry. cDC=conventional dendritic cells; pre-cDC=pre-conventional dendritic cells; Neut.=neutrophils; NK cell=natural killer cells; infDC=inflammatory dendritic cells; MDSC=myeloid-derived suppressor cells; pDC=plasmacytoid dendritic cells; pre-pDC=pre-plasmacytoid dendritic cells.
